# Sequence architecture shapes human allele frequencies through ectopic gene conversion

**DOI:** 10.64898/2026.08.26.747289

**Authors:** Wouter Steyaert

## Abstract

Population-genetic interpretations of allele frequency commonly begin from a single-origin assumption: that a variant arose once and was subsequently shaped by selection, drift and demographic history. Here we show that for a substantial fraction of human variation this assumption fails, and fails predictably: ectopic gene conversion reintroduces the same nucleotide change generation after generation, at rates strongly structured by two fixed properties of genome architecture, homologous template length and donor-acceptor distance. Across 534 million gnomAD v4.1 variants, this recurrent input accounts for an estimated 4% of the rarest variants and more than 15% of common ones. De novo mutations in 11,963 trios show the process directly: the mutation rate is elevated up to 28-fold precisely where the same variant is already common, with the newly arising allele matching the paralogous donor in 94% of such cases at long templates. The effect is strongest within segmental duplications but extends along every chromosome. Affected positions show reduced linkage disequilibrium and, at common frequencies, are depleted by 30–40% among reported GWAS associations. For this fraction of human variation, allele frequency is encoded in genome sequence architecture rather than set by population-genetic processes alone.

## Introduction

The infinite-sites assumption treats each variant as having arisen only once in the population^1^. It underpins classical treatments of the site frequency spectrum and coalescent- and haplotype-based inference, in which recurrent mutation is neglected and allele frequencies are interpreted primarily as reflecting selection, drift, and demographic history. Yet ectopic gene conversion violates this assumption systematically. During meiosis, double-strand break (DSB) repair^2^ can use a paralogous sequence rather than the homologous chromosome as template, copying nucleotide differences from donor to acceptor in a non-reciprocal transfer^3,4^. Because the same paralogous donor can serve as template in independent meioses, the same donor-specified change can recur at the acceptor position across generations. If such recurrence is widespread, its rate becomes an additional determinant of allele frequency alongside selection, drift, and demographic history.

Ectopic gene conversion is well established. Conversion between paralogs drives concerted evolution in gene families^5^, and conversion from pseudogenes can introduce pathogenic variants into protein-coding genes at loci such as *CYP21A2*^3^. But existing evidence has largely come from individual loci and gene families. Recent work estimated ectopic conversion between gene duplicates at approximately 20 times the point mutation rate^6^, but its effect on standing variation has not been measured genome-wide. Non-crossover meiotic gene conversion between homologous chromosomes has been quantified at approximately 5.9–7.0 × 10^-6^ per base pair per generation in humans^7,8^, but no comparable estimate exists for ectopic conversion between paralogs. Without one, it remains unclear whether ectopic gene conversion is largely confined to duplicated regions or is a major determinant of the allele frequency spectrum.

Genome-wide evidence has focused largely on segmental duplications (SDs). These regions, >1 kb in length with >90% sequence identity, comprise ∼5% of the genome^9^. Vollger et al.^10^ showed that single-nucleotide variant (SNV) density is elevated by 60% in SDs, with at least 23% of this excess attributable to ectopic gene conversion. This is consistent with the earlier identification of ∼16,000 variants matching their paralogous donor in these regions^11^ and is further supported by long-read sequencing of families, which has confirmed increased mutation rates in repetitive DNA, with SD mutability dependent on length and sequence identity^12^. But SDs represent only a fraction of the paralogous homology in the genome; shorter stretches of near-identical sequence are far more numerous, and their contribution has not been quantified.

Here we map all positions in the human genome where a paralogous template can introduce a specific nucleotide change and analyse 534 million gnomAD v4.1^13^ variants together with de novo mutations (DNMs) from 11,963 trios. We show that the rate of ectopic gene conversion scales with two architectural parameters, rising with homologous template length and falling with donor-acceptor distance. This recurrent mutational input is a major determinant of allele frequency for a substantial fraction of human variation, and its effect is not confined to SDs but extends along the full length of every chromosome.

## Results

### Gene conversion-compatible positions are enriched for genetic variants

We identified all genomic positions where ectopic gene conversion could introduce an SNV by mapping k-mer pairs (k = 17–91 bp) that differ only at their central nucleotide (Figure 1a–c; Methods). At these conversion-compatible positions, the repair machinery could replace the central nucleotide of the acceptor with that of the donor. The number of such positions ranges from 1.7 million for k = 91 to 2.1 billion for k = 17, corresponding to 0.06% and 79.8% of the mappable genome, respectively (Supplementary Table 1).

**Figure 1:**
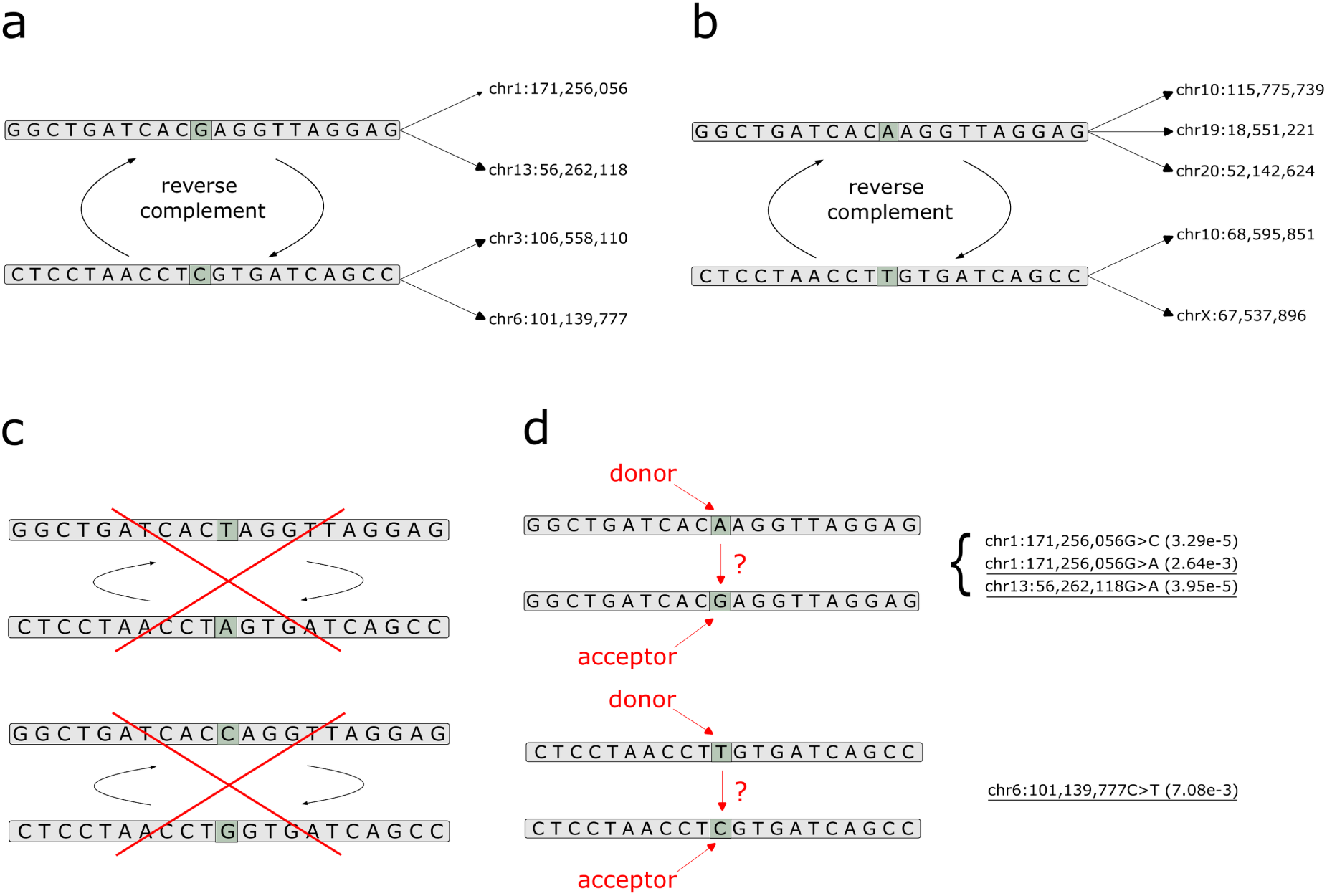
Identification of gene conversion-compatible positions. (a) A 21-mer with central nucleotide G occurs at four genomic positions (two on the forward strand, two as reverse complement). (b) A 21-mer identical except for central nucleotide A occurs at five additional positions. The central positions in (a) and (b) are gene conversion-compatible: conversion between paired sequences would introduce a G↔A change. (c) The remaining possible central nucleotides (T and C) do not form valid k-mer pairs, as no matching sequence exists elsewhere in the genome; these positions are not conversion-compatible. (d) Gene conversion from donor to acceptor would replace the central nucleotide of the acceptor with that of the donor. At the conversion-compatible positions in this example, four variants are present in gnomAD v4.1 (allele frequencies in parentheses). Three are concordant with the predicted donor-to-acceptor change (underlined); one is discordant (not underlined). An excess of concordant over discordant variants is the signature of ectopic gene conversion.

Using 534 million variants from gnomAD v4.1, we asked at each conversion-compatible position whether a variant is present and, if so, whether the change matches the predicted donor-to-acceptor transfer (Figure 1d). An excess of such “concordant” variants over discordant ones provides a nucleotide-specific signature of gene conversion.

Variants are enriched at conversion-compatible positions across all combinations of template length and allele frequency (AF; Figure 2a; all Bonferroni-corrected p < 10^-17^; Supplementary Tables 2, 3). The magnitude depends strongly on template length: for k = 91, variant density exceeds expectation by more than 450% in the highest frequency class (AF > 0.5), and by 325–454% across all common variant bins (AF ≥ 0.05); for k = 17, the excess is only 1–2%. Because short homologous sequences vastly outnumber long ones, their collective contribution exceeds that of longer templates despite the weaker per-position effect.

**Figure 2:**
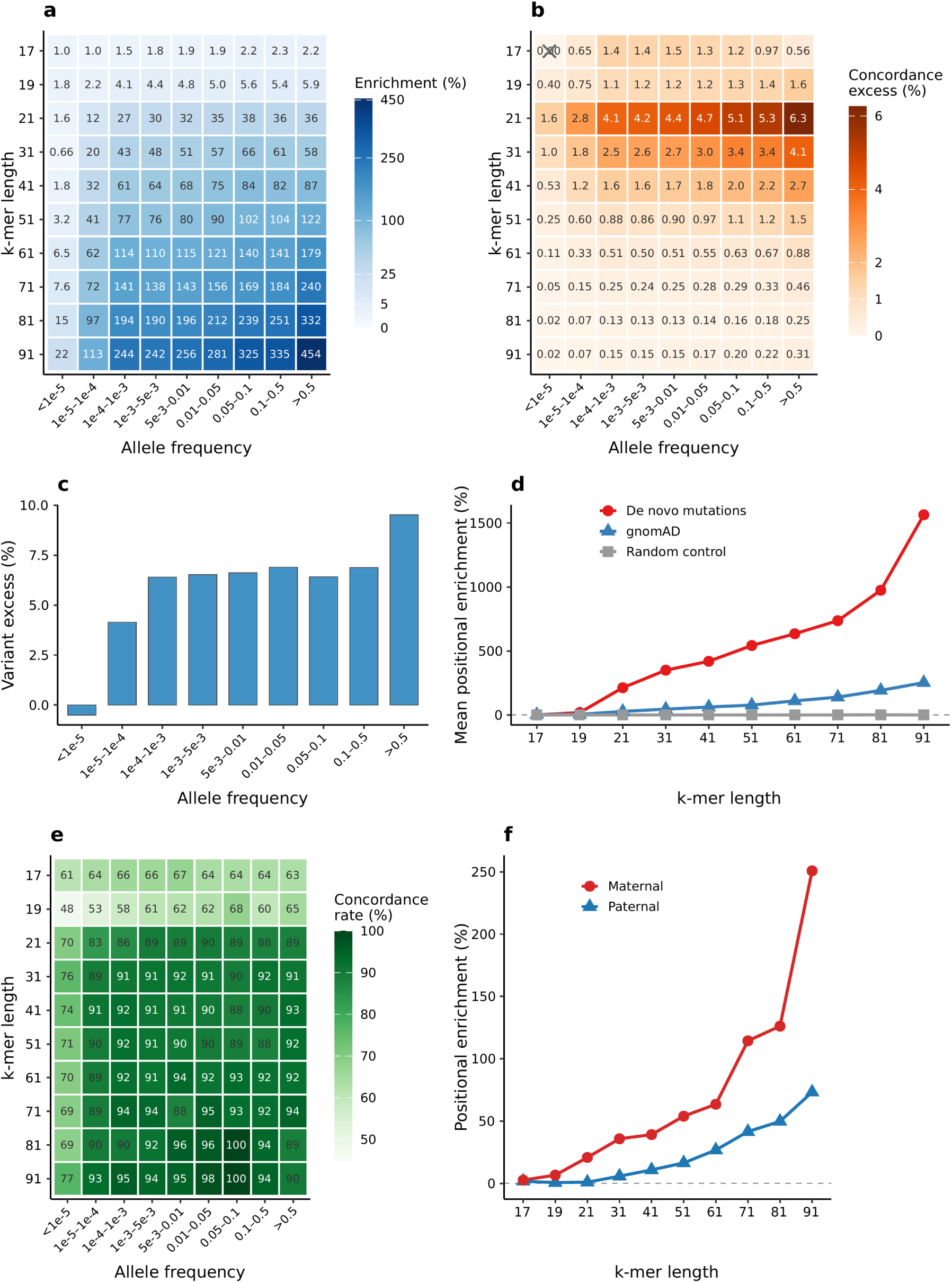
Variant enrichment at gene conversion-compatible positions. (a) Relative enrichment of variant density at conversion-compatible versus non-compatible positions, shown for each combination of template length k (rows) and gnomAD v4.1 allele frequency bin (columns). (b) Concordance excess: the percentage of all genome-wide variants in each frequency class attributable to gene conversion. Concordance excess decreases with increasing k because longer templates occupy progressively fewer genomic positions, reducing their cumulative contribution to total variation despite stronger per-position enrichment (panel a). The crossed cell (k = 17, allele frequency < 10^−5^) indicates the only non-significant result (Bonferroni-corrected p > 0.05). (c) Alignment-based validation: excess of gnomAD variants over randomly placed variants harbouring a BLAST-detected homologous tract of at least 21 bp on the reference genome with the alternative allele at the central position, per allele frequency bin. (d) Mean positional enrichment averaged across allele frequency bins as a function of k-mer length for de novo mutations (red), gnomAD population variants (blue), and randomly placed variants (grey). (e) Concordance rate of de novo mutations with the predicted donor allele. (f) Positional enrichment of maternally (red) versus paternally (blue) derived de novo mutations at positions with rare population variants (allele frequency < 10^−5^).

At conversion-compatible positions, concordant variants show extreme enrichment over expectation, strongly implicating gene conversion rather than a non-specific property of these genomic regions (Figure 2b; Bonferroni-corrected p < 10^-14^ for 89 of 90 tests; Supplementary Tables 2, 3). The single exception is k = 17 at the lowest frequency class (AF < 10^-5^).

The enrichment increases markedly with AF: the variant excess is approximately 1% among the rarest variants (AF < 10^-5^), rising to 11% among common variants (AF ≥ 0.05) (Methods; Supplementary Table 2). This frequency dependence is expected under heterogeneous recurrent mutation rates: variants at higher-rate positions are repeatedly reintroduced and therefore become increasingly represented at higher frequencies, whereas variants at lower-rate positions remain predominantly rare under drift. The concordance signal attributes an even larger fraction of variants to gene conversion: 4% of the rarest, more than 15% of common variants (AF ≥ 0.05), and 19% at the highest frequencies (AF ≥ 0.5) (Figure 2b; Supplementary Table 2). The gap between the concordance signal and the positional excess is itself consistent with recurrence: it reflects a depletion of discordant variants below the random-mutation baseline, as expected under ongoing replacement by the donor-matching allele, whereas a one-time event would contribute equally to both signals.

Alignment-based homologous tract detection around randomly sampled gnomAD variants (Methods) independently reproduced the frequency dependence, from no detectable excess at the rarest AFs (−0.5%, indistinguishable from baseline) to 9.5% at AF > 0.5 (Supplementary Table 4; Figure 2c). As a second validation, 10 million simulated single-nucleotide substitutions placed uniformly at random across the genome showed no enrichment at conversion-compatible positions for any k (Supplementary Table 5). A third, technology-orthogonal validation recomputed variant-density enrichment using PacBio HiFi long-read data (Genomic Answers for Kids^14^, n = 541; Methods). HiFi-based enrichment reproduced the gnomAD-based enrichment across all strata and template lengths (ratio 0.96–1.30). Within segmental duplications at k ≥ 51, HiFi enrichment is comparable to or greater than the gnomAD value, opposite to the reduction expected if mismapping had inflated the gnomAD calls (Supplementary Table 6; Supplementary Figure 1). Variants at conversion-compatible positions were confirmed in HiFi at 94–99% (AF ≥ 0.01; Supplementary Table 7), bounding any mismapping-driven inflation of the enrichment to at most 1.009× (Supplementary Table 8; Supplementary Figure 2).

From templates of 21 bp upward, enrichment at conversion-compatible positions is consistently stronger within SDs than outside (Supplementary Table 9). For k = 91 and variants with AF above 0.1, variant density is elevated by at least 805% within SDs versus 301% outside. Yet the effect outside SDs remains substantial: the cumulative excess accounts for 12.1% of common variants (AF 0.1–0.5; Supplementary Table 9b), and even for shorter templates (k = 21–41), variant density is elevated by 34–82%.

### De novo mutations confirm recurrent mutation and support a meiotic origin

DNMs provide a direct test of ongoing recurrence. If ectopic gene conversion drives recurrent mutation, the DNM rate at compatible positions should be highest for nucleotide changes that are already common in the population. We examined 865,392 DNMs from 11,963 parent-offspring trios^15^ (Methods).

DNMs are enriched at conversion-compatible positions, and the enrichment increases sharply with both template length and population AF (Figure 2d; Supplementary Tables 10 and 11). For k = 91, the DNM rate is elevated by 97% above the non-compatible background at positions with rare population variants (AF < 10^-5^) but by 2,731% where the same variant is common (AF 0.1–0.5; 177 observed versus 6.25 expected, equivalently ∼28-fold; Bonferroni-corrected p < 10^-325^). For k = 21, the corresponding values are 8% and 296%. Even for k = 17, where nearly 80% of the genome is compatible, a modest but significant enrichment persists at the rarest variants (2.5% at AF < 10^-5^; Bonferroni-corrected p = 1.3 × 10^-11^). Under a single-origin model, the population frequency of a variant should not predict the rate at which the same nucleotide change arises de novo.

The observed relationship demonstrates directly that the same changes arise repeatedly and links the rate of recurrence to population AF.

The identities of the nucleotide changes provide a second mechanistic test. Among DNMs at conversion-compatible positions with k ≥ 21, concordance with the predicted donor allele is 70% at positions with rare population variants and rises to 94% at positions with common variants and long templates (Figure 2e; Supplementary Table 10). At the longest templates and highest frequencies, DNMs almost always introduce the exact nucleotide predicted by gene conversion.

Stratification by parent of origin reveals a signature consistent with meiotic origin. Paternally derived DNMs outnumber maternally derived DNMs 3.4 to 1 among the 309,026 successfully phased autosomal variants (Methods), consistent with the greater number of mitotic cell divisions in spermatogenesis compared with oogenesis^16^. Yet at positions with rare population variants (AF < 10^-5^), maternally derived DNMs show consistently stronger enrichment at conversion-compatible positions: at k = 91, maternal enrichment is 251% compared with 73% paternal, a more than three-fold difference (Figure 2f; Supplementary Table 12). This asymmetry diminishes with increasing AF and vanishes at common variants (AF > 0.1), where paternal and maternal enrichment converge (both exceeding 3,000%).

This is the pattern expected if ectopic gene conversion is predominantly meiotically driven, with the paternal signal diluted by a large excess of mitotically derived DNMs at random positions (see Discussion).

These enrichment data allow an empirical estimate of the per-generation ectopic gene conversion rate. For each template length, the excess of concordant DNMs over the discordant background, divided by the number of trios (N = 11,963) and conversion-compatible positions, yields r_gc, the per-site per-generation rate of ectopic conversion-driven allele change (Methods; Supplementary Table 13). Across all mappable regions, r_gc increases from 7.6 × 10^-11^ at k = 17 to 1.4 × 10^-8^ at k = 91, corresponding to 0.01× and 2.1× μ_eff, the discordant de novo mutation rate estimated locally at the same conversion-compatible positions (Methods), respectively. Within SDs, the rate rises steeply: at k = 91, r_gc reaches 8.1 × 10^-8^ (95% CI: 6.4–9.8 × 10^-8^), corresponding to 15.4-fold (95% CI: 7.0–32.7) μ_eff (see Discussion). Outside SDs, the rate at k = 91 still reaches 1.1 × 10^-8^ (1.7× μ_eff), and at shorter templates (k = 21) drops to 0.4× μ_eff, confirming that ectopic conversion operates genome-wide, albeit at reduced efficiency.

### Enrichment varies across the genome

Across non-overlapping 1-Mb windows spanning mappable regions of all chromosomes, the great majority show positive enrichment at conversion-compatible positions (Figure 3a; Supplementary Figure 3; Methods). The most strongly enriched windows cluster into 193 distinct genomic regions (Figure 3b; Supplementary Table 14; Methods). The majority of these regions are interstitial (64%), with the remainder pericentromeric (26%) and subtelomeric (10%). Although 83% overlap annotated SDs, the predominance of interstitial regions shows that strong enrichment is not restricted to the pericentromeric and subtelomeric regions where SDs are most concentrated.

**Figure 3:**
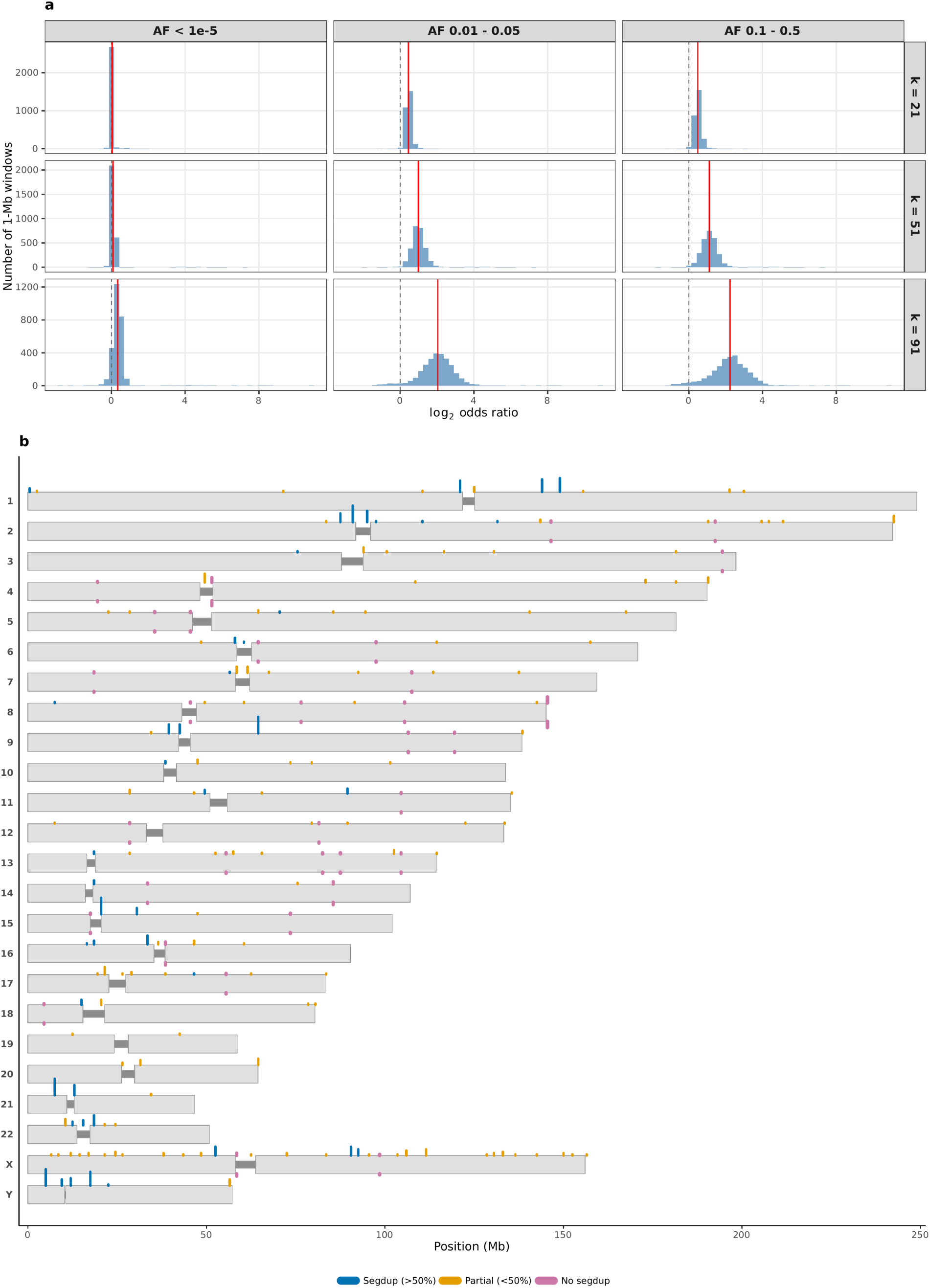
Genome-wide enrichment at gene conversion-compatible positions. (a) Distribution of log_2_ odds ratios across non-overlapping 1-Mb windows, shown as histograms faceted by template length k (rows; k = 21, 51, 91) and allele frequency bin (columns; AF < 10^−5^, 0.01–0.05, 0.1–0.5). Grey dashed vertical lines mark log_2_(OR) = 0 (null expectation); red solid vertical lines mark the per-panel median. (b) Ideogram showing the locations of 529 within-stratum peak calls (across all template-length × allele-frequency strata; corresponding to 193 distinct genomic regions) across all chromosomes, coloured by segmental duplication overlap: blue = > 50% segdup, orange = partial overlap (< 50%), pink = no segdup overlap. The grey bands within each chromosome indicate the centromere.

Thirty-three of these regions (17%) fall entirely outside SDs. Among these, a peak at chr11 (104 Mb; k = 21) harbours the tandem caspase paralogs *CASP4*, *CASP5* and *CASP12*, and a peak at chr7 (107 Mb; k = 21) contains the solute carrier paralogs *SLC26A3* and *SLC26A4*. Such tandem families provide multiple paralogous templates within a short interval, consistent with strong enrichment outside annotated SDs. After excluding all peak windows and recomputing the genome-wide enrichment test, enrichment at conversion-compatible positions remained significant for every combination of template length and allele frequency (all Bonferroni-corrected p < 10^-15^, m = 90; Supplementary Table 15). Thus, ectopic gene conversion is concentrated in hotspots but remains significant across the rest of the genome.

### Recombination hotspots show elevated ectopic conversion

If ectopic gene conversion is initiated by meiotic DSBs, enrichment at conversion-compatible positions should be elevated within recombination hotspots, where meiotic recombination is concentrated. From k = 21 onward, enrichment is higher inside than outside hotspots at every template length, and the difference increases steeply with k, reaching approximately 1% at k = 21, 7% at k = 31 and 88% at k = 91 (Supplementary Table 16; Methods). At k = 91, all nine AF bins show significantly higher enrichment inside than outside hotspots (Woolf heterogeneity test, Bonferroni-corrected p < 0.05). At k = 17 and k = 19, the inside-versus-outside difference is negligible (<1%).

Mean concordance with predicted donor alleles inside hotspots follows the same template-length dependence, rising from 76% at k = 21 to 81–84% for k ≥ 31, while remaining markedly lower at k = 17 and k = 19. At longer templates, the hotspot-associated excess is predominantly donor-concordant, supporting meiotic ectopic conversion rather than a non-specific increase in hotspot-associated mutation rate.

### Conversion signal decays with donor-acceptor distance

Beyond template length, the physical distance between donor and acceptor should independently modulate ectopic conversion: strand invasion is expected to be favoured by spatial proximity during meiosis. We restricted the analysis to unambiguous one-to-one donor-acceptor pairs, stratified by genomic distance into seven intra-chromosomal bins and one inter-chromosomal category, and computed the mean AF of conversion-consistent variants in each bin (Methods; Supplementary Table 17).

Mean AF of conversion-consistent variants decreases steeply with donor-acceptor distance. At k = 21, mean AF drops from 4.5% for pairs separated by less than 1 kb to 0.7% at distances exceeding 100 Mb, a 6.5-fold reduction (Figure 4a). The decay is steepest between 100 kb and 10 Mb, with AF approaching inter-chromosomal levels beyond 10 Mb, indicating that linear distance becomes a weak predictor of the conversion signal beyond the megabase scale. The overall distance dependence is observed across template lengths, while increasing k amplifies the signal: at the shortest distances (<1 kb), mean AF rises from 3.8% at k = 17 to 9.6% at k = 81. At k ≥ 61 and distances below 1 kb, more than 65% of donor-acceptor pairs carry a conversion-consistent gnomAD variant, consistent with nearby, highly identical paralogues being efficient substrates for recurrent conversion. At the longest template length analysed (k = 91), the highest mean AF occurs not in the closest distance bin (<1 kb, AF = 9.2%) but at 10–100 kb (AF = 11.4%). Because the 91-bp criterion does not capture the true tract length underlying each pair, this may reflect longer tracts on average at intermediate distances rather than a distance effect per se. Intra-chromosomal pairs show consistently higher AF than inter-chromosomal pairs at every template length (Figure 4b), with the gap widening from 2.0-fold at k = 21 to ∼2.4-fold at k = 61 before returning towards 2.0-fold at k = 91. Inter-chromosomal pairs nonetheless retain a detectable conversion signal (mean AF = 3.6% at k = 91).

**Figure 4:**
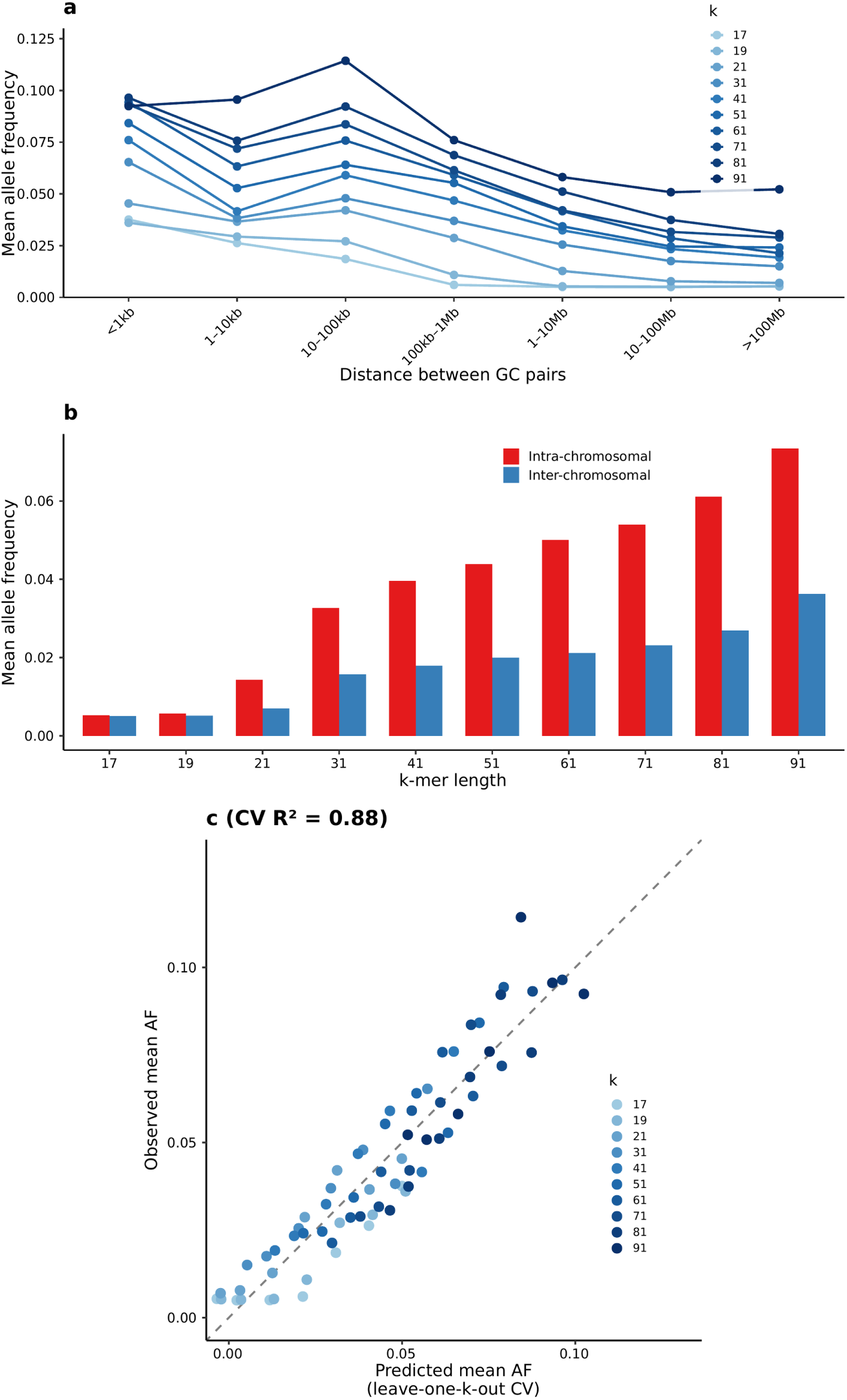
Allele frequency is determined by sequence architecture. (a) Mean allele frequency as a function of donor-acceptor distance for each template length. (b) Mean allele frequency for intra-chromosomal versus inter-chromosomal donor-acceptor pairs as a function of template length. (c) Predicted versus observed mean allele frequency for each k × distance combination, based on leave-one-k-out cross-validation. Points are coloured by template length k.

Together, template length and donor-acceptor distance strongly predict mean AF across architectural strata. A simple regression on these two parameters, validated by leave-one-k-out cross-validation, explains 88% of the variance in observed bin-mean AF across all k × distance combinations (R^2^ = 0.88; Figure 4c). Both parameters are fixed properties of genome sequence architecture, showing that the frequency distribution of conversion-consistent variants is strongly structured by predictable recurrent mutational input.

### Ectopic conversion shows a GC-biased signature

The AF signal also carries the expected signature of GC-biased resolution of heteroduplex intermediates. The ratio of mean AF for W→S versus S→W variants, where W denotes A/T and S denotes G/C, increases from 1.2× at k = 17 to 3.3× at k = 61 (Supplementary Figure 4). The W→S fraction rises from ∼30% at low AFs, consistent with the known ∼2-fold excess of S→W mutation, to above 50% at AF > 0.5 for k ≥ 31 and at AF > 0.01 for k ≥ 81 (Supplementary Table 18).

This shift from S→W predominance at low frequencies to W→S enrichment at higher frequencies is consistent with GC-biased mismatch repair during heteroduplex resolution, a mechanism established for allelic gene conversion^8,17^ and here observed as a population-level signature of conversion between paralogs.

### Reduced linkage disequilibrium and depletion of GWAS associations at conversion-compatible positions

Recurrent introduction of the same allele onto independent haplotype backgrounds should weaken its association with any single set of surrounding markers, predicting lower linkage disequilibrium (LD) than for a single-origin variant^18^. We tested this in phased 1000 Genomes high-coverage genotypes^19^, comparing mean r^2^ between variants at conversion-compatible positions and frequency-matched variants at non-compatible positions across template length (k) and AF bins (Methods; Supplementary Table 19). Across template lengths and AF bins, mean r^2^ is lower at conversion-compatible than at matched non-compatible positions, with the few exceptions confined to low-count strata (Figure 5a, k = 21, 41 and 71; all 10 k values in Supplementary Table 19). The reduction becomes stronger at longer k, reaching 46% in the 1–5% AF bin at k = 31 (mean r^2^ 0.074 versus 0.135; Wilcoxon p = 4.3 × 10^-9^) and 64% at k = 51 (p = 2.6 × 10^-10^), and persists at common AFs (20% reduction at k = 31, AF 0.1–0.5; p = 1.5 × 10^-3^). The pattern is robust across European and all 1000 Genomes samples, window sizes (10–100 kb) and local recombination-rate strata (Supplementary Tables 19, 20), arguing against local recombination as the primary explanation.

**Figure 5:**
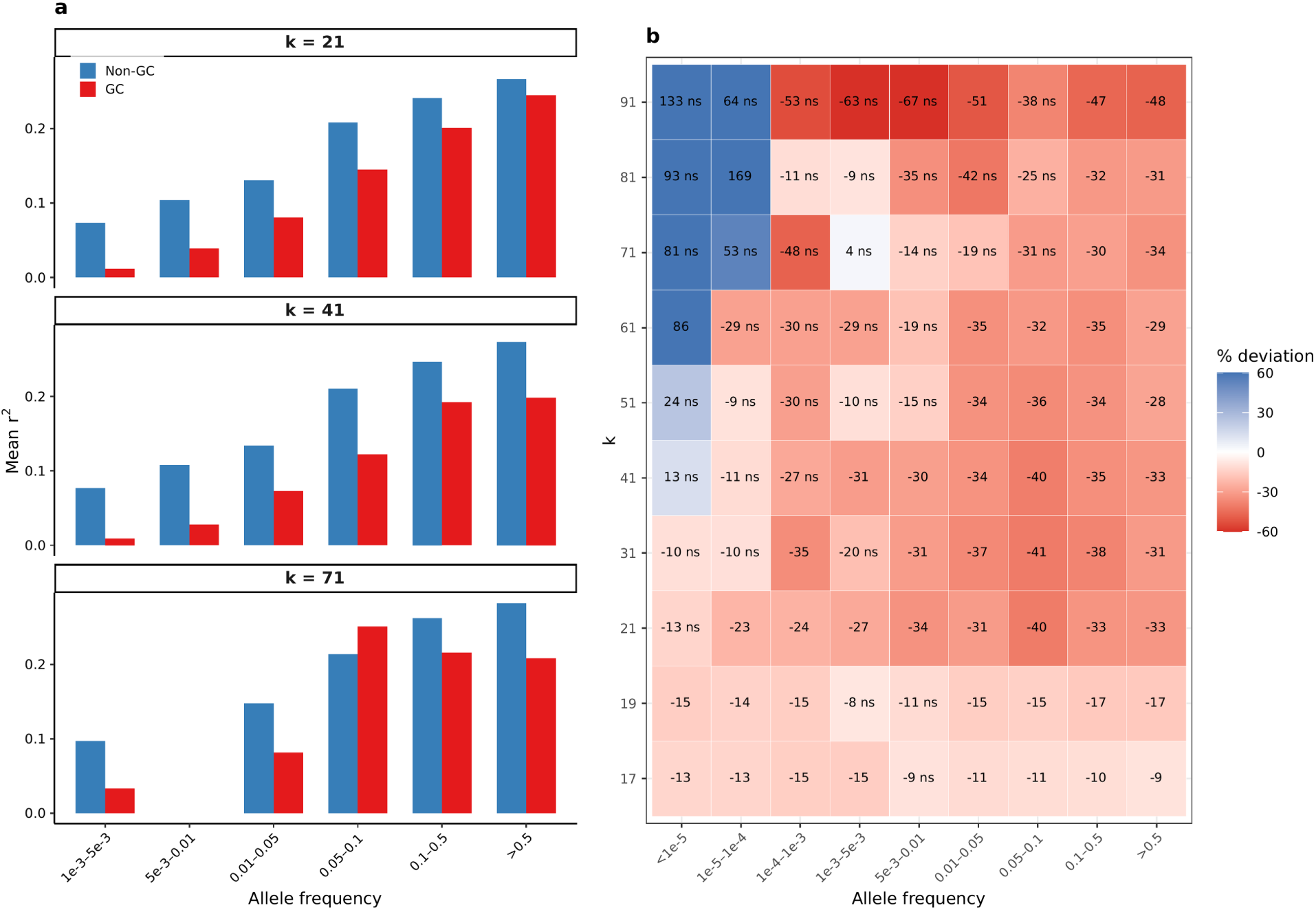
Linkage disequilibrium and GWAS depletion at conversion-compatible variants. (a) Cell-mean r^2^ between conversion-compatible (GC, red) and non-compatible (Non-GC, blue) variants across matched (k × allele frequency) cells, shown for k = 21, 41 and 71 (EUR, 1000 Genomes high-coverage, 10 kb window, 1:1 paralog pairs; Supplementary Table 19). Conversion-compatible positions show systematically lower r^2^ than frequency-matched non-compatible positions, with a single exception at k = 71, allele frequency 0.05–0.1 (n = 34), consistent with recurrent mutation as a major contributor to reduced LD at these sites. The k = 71 AF 5×10^−3^–10^−2^ cell is dropped (both GC and Non-GC bars omitted) because the GC variant count fell below the n = 30 threshold after filtering. (b) Depletion of GWAS Catalog variants at conversion-compatible positions versus matched non-compatible positions, expressed as percentage difference in variant density [(GC freq − non-GC freq) / non-GC freq × 100], shown as a heatmap over all k × allele-frequency cells (All mappable regions; Supplementary Table 21). Negative values (red) denote depletion; positive values (blue) denote enrichment. Cells annotated with ‘ns’ indicate non-significant deviations (Bonferroni-corrected p > 0.05) and correspond to low-count strata with as few as 4 GWAS variants per cell. Depletion is consistent and significant across k ≥ 21 at common allele frequencies (AF ≥ 0.05), plateauing in the 30–40% range; most blue cells at long k and rare AF are non-significant, with two exceptions that remain significant despite low counts (k = 81, AF 10^−5^–10^−4^; k = 61, AF < 10^−5^).

Lower LD could reduce how efficiently common variants at conversion-compatible positions are tagged by nearby markers and therefore their probability of appearing as GWAS associations. We compared the density of GWAS Catalog associations^20^ at conversion-compatible versus non-compatible positions across the same k × AF grid. GWAS-associated variants are depleted at conversion-compatible positions, with the few exceptions confined to low-count strata at long templates and rare AFs (Figure 5b; Supplementary Table 21). The depletion increases with template length and AF, reaching approximately 30–40% across k = 21–41 at common frequencies: 33%, 38% and 35% at k = 21, 31 and 41, respectively, for AF 0.1–0.5 (Bonferroni-corrected log10 p = −323, −190 and −92). The pattern is consistent across mappability strata, including within and outside SDs (Supplementary Table 21).

## Discussion

The infinite-sites model treats each segregating variant as the product of a single mutational event. Our results show that this approximation fails systematically for a substantial fraction of human variants, from 4% of the rarest (AF < 10^-5^) to more than 15% of common variants (AF ≥ 0.05). The frequency dependence of this fraction is consistent with mutation-rate heterogeneity across positions, with recurrent mutational input disproportionately contributing to variants at higher frequencies. Ectopic gene conversion provides this recurrent input: the frequencies of affected variants are shaped not only by selection, drift and demographic history, but also by the rate at which sequence architecture reintroduces the same change generation after generation. This recurrence shifts the site frequency spectrum toward common variants^21^ and reduces LD; affected positions are also depleted among reported GWAS associations. Methods that ignore this heterogeneity could consequently bias demographic and selection inference, potentially generating patterns interpreted as population expansion or balancing selection.

Previous studies focused on SDs, where long homologous tracts create favourable conditions for ectopic gene conversion^10^. Consistent with this, enrichment is markedly stronger within SDs than outside, but the signal extends continuously along entire chromosomes, well beyond annotated duplications into the broader repeat landscape. The cumulative contribution of shorter homologous sequences (k = 17–41 bp), which pervade the genome, exceeds that of longer templates despite their weaker per-position effects.

The rate varies by orders of magnitude across the genome, from negligible where no suitable donor exists to above the point mutation rate at long, nearby homologies. Within SDs at k = 91, the ectopic conversion rate is 15.4-fold μ_eff, the locally estimated discordant DNM rate (r_gc = 8.1 × 10^-8^; 95% CI: 6.4–9.8 × 10^-8^), comparable in order of magnitude to the 2.5 × 10^-7^ per generation estimated by Harpak et al.^6^ from evolutionary divergence between gene duplicates, an approximately 20-fold excess over the point mutation rate. The convergence is noteworthy because the two estimates are methodologically independent, although not identically normalised: our fold excess is relative to μ_eff, a locally estimated discordant DNM rate that spans two of the three possible alternative alleles at each position, whereas theirs is relative to a per-base-pair rate. Our normalisation is therefore conservative — expressing the excess relative to a single-allele rate would raise it by up to approximately two-fold — and the two estimates are of the same order rather than directly equivalent. Outside SDs, the rate at the same template length drops to 1.7-fold, and at shorter templates (k = 21) to 0.4-fold, establishing a spectrum from rare conversion events at short homologies to rates exceeding point mutation at long ones. This spectrum is distinct from allelic (non-crossover) gene conversion, which proceeds at approximately 5.9–7.0 × 10^-6^ per base pair per generation^7,8^, roughly two orders of magnitude above the ectopic rate within SDs at k = 91. Allelic conversion acts on heterozygous sites between homologous chromosomes rather than on fixed paralogous differences and therefore does not provide the same mechanism for recurrent introduction of donor-specified mutations across generations.

Two converging lines of evidence point to a predominantly meiotic rather than replication-associated origin. Enrichment at conversion-compatible positions is elevated within meiotic recombination hotspots, where meiotic DSBs concentrate. The parent-of-origin asymmetry in DNMs, with stronger maternal enrichment at rare-variant positions and convergence at common variants, is consistent with meiotically derived events contributing proportionally more to the maternal DNM set, while the paternal signal is diluted by the larger contribution of mitotic events. Replication-associated mechanisms such as template switching would not predict concentration at meiotic recombination hotspots. Together with the observed 70–94% concordance of DNMs with the predicted donor allele, this supports DSB-mediated ectopic conversion as the underlying process. The GC bias in AF, with W→S variants reaching higher frequencies than S→W variants, is also consistent with biased mismatch repair during heteroduplex resolution, the same mechanism established for allelic gene conversion^8,17^.

Concordance with the donor allele could also arise in part from nucleotide context rather than conversion. Paralogous differences have themselves arisen by mutation and are enriched for transitions and hypermutable CpG dinucleotides, so independent mutations may match the donor allele more often than expected under a context-free model. Three architectural dependencies argue against nucleotide context as the primary explanation. Enrichment scales strongly with template length, from 1–2% at k = 17 to more than 450% at k = 91, yet whether a position is a CpG does not depend on the length of the surrounding homology. Mean allele frequency declines with donor-acceptor distance and approaches inter-chromosomal levels at large separations, a dependence for which local nucleotide context provides no obvious explanation. Hotspot enrichment is likewise not explained by nucleotide context alone, because local base composition does not predict why the donor-concordant excess should strengthen specifically within meiotic recombination hotspots. Nucleotide context may therefore modulate absolute mutation rates, but it cannot account for the combined dependencies on template length, donor-acceptor distance and meiotic recombination.

Reduced LD at conversion-compatible positions is consistent with recurrent mutation introducing the same allele onto independent haplotype backgrounds and thereby weakening its association with surrounding markers. Common variants at these positions are also depleted among reported GWAS associations by approximately 30–40%, consistent with less efficient tagging by nearby variants. This association does not by itself establish reduced LD as the sole cause of GWAS depletion, but it identifies a potential consequence of recurrent mutational origin for association mapping. The same LD reduction could also affect haplotype-based tests of natural selection that assume common alleles share a single origin on an extended haplotypic background, potentially mimicking signatures of balancing selection or obscuring genuine selective sweeps at affected sites.

Ectopic gene conversion may also have therapeutic implications. CRISPR/Cas9-induced DSBs can be repaired using a paralogous sequence as an endogenous template, correcting pathogenic mutations without exogenous donor DNA, as demonstrated at up to 27% efficiency for the β0 41/42 mutation in *HBB* using *HBD* as the endogenous template^22^. Our map may therefore help identify loci at which endogenous paralogous templates could be exploited for DSB-induced repair and prioritize candidate sites according to architectural features associated with conversion. Natural germline conversion rates need not translate directly into editing efficiency, however. Conversely, the map highlights positions at which DSB-inducing edits may warrant consideration of unintended conversion from paralogous donors as a potential editing outcome.

Our estimates are likely conservative. Templates outside the analysed range may contribute additional signal. The analysis is restricted to SNVs treated as independent events, whereas a single conversion tract can introduce short indels, multi-nucleotide changes or several linked SNVs, all of which are excluded from the present estimates and could increase the fraction of variation attributable to this process. Substantial variation in enrichment among positions with identical template length further indicates that additional factors, including chromatin context, flanking sequence and many-to-many donor-acceptor relationships, modulate conversion efficiency beyond template length and distance.

Mutation-rate variation is a primary determinant of allele-frequency distributions^21,23^. Ectopic gene conversion is one source of such variation, but one whose rate is systematically structured by genome architecture. It rises with homologous template length and declines with donor-acceptor distance, two properties that can be read directly from the reference genome (Figure 4c). For more than 15% of common human variants, allele frequency is therefore not a readout of selection, drift and demographic history alone; it is also partly encoded in genome sequence architecture through predictable recurrent mutational input.

## Methods

### Identification of gene conversion-compatible positions

We scanned the GRCh38 human reference genome for positions susceptible to ectopic gene conversion. For each template length k in {17, 19, 21, 31, 41, 51, 61, 71, 81, 91}, we extracted all k-mers from the chromosomal FASTA sequences using a sliding window of one nucleotide, recording both forward and reverse-complement orientations along with their genomic coordinates. K-mer occurrences were stored per chromosomal chunk as gzipped text files keyed by the k-mer string, then aggregated genome-wide per 5-nucleotide seed (the first five nucleotides of the k-mer) into per-seed summary files.

A position was defined as gene conversion-compatible if at least one pair of k-mers exists in the genome that are identical except at their central nucleotide (position (k−1)/2 + 1). For each compatible position, we recorded the reference allele, the observed donor allele(s) (the central nucleotide(s) of matching k-mers, with appropriate complementation for reverse-complement matches), and the recurrence count (number of distinct donor templates supporting that change). Donor coordinates are retained in the upstream per-seed coordinate files but are not propagated to the per-position BED. The central nucleotide was identified by extracting the flanking (k−1)/2 bases on each side, then searching for k-mers sharing these flanks but carrying a different central base; both the forward sequence and its reverse complement were considered.

Positions compatible at multiple template lengths were assigned exclusively to the longest compatible length by computing set differences between consecutive k values: for k < 21, consecutive values differ by 2 (e.g. 17 vs 19); for k ≥ 21, by 10 (e.g. 21 vs 31). Specifically, the set of positions exclusive to template length k was defined as all positions compatible at k but not at k+2 (for k < 21) or k+10 (for k ≥ 21). This was implemented by computing per-chromosome difference files (gcdiffs) representing positions uniquely compatible with each k interval; for k = 91, the full set of compatible positions was used directly. Gene conversion files for each k-mer length were constructed by merging these difference files with the next-longer k conversion file. Per-chromosome BED files containing both positional and nucleotide-change information were then sorted, bgzipped, and tabix-indexed for efficient downstream retrieval, supporting both positional and concordance-based tests. The longest category (k = 91) contains all positions with compatible templates ≥ 91 bp.

Genome-wide coverage of conversion-compatible positions was tabulated against the GIAB v3.5 all-mappable mask (2.69 Gb); the mappability stratification used in downstream contingency tests is described under Enrichment analysis.

### Variant data

SNVs were obtained from gnomAD v4.1 genomes^13^ (gnomad.genomes.v4.1.sites.chrN.vcf.bgz). Only PASS-filtered single-nucleotide substitutions were retained (single-base reference and alternative alleles). Allele frequency (AF), allele count (AC), and allele number (AN) were parsed from the VCF INFO field. Variants were stored in sorted, tabix-indexed BED format per chromosome, concatenated across chromosomes, and restricted to the GIAB v3.5 all-mappable mask using bedtools intersect; this final set of 534 million SNVs is the input for all downstream analyses. For the enrichment analyses (see Enrichment analysis) variants were stratified into nine AF bins: < 10^−5^, 10^−5^ to 10^−4^, 10^−4^ to 10^−3^, 10^−3^ to 5×10^−3^, 5×10^−3^ to 10^−2^, 10^−2^ to 5×10^−2^, 5×10^−2^ to 0.1, 0.1 to 0.5, and ≥ 0.5.

### De novo mutation data and processing

DNMs were obtained from the Genomics England 100,000 Genomes Project^15^ denovo variant dataset (LabKey V9). De novo variant VCF files were processed per trio. Variant-level filtering removed calls flagged by the Genomics England DNM caller as base_fail, altreadparent, or abratio, and retained only single-nucleotide substitutions with a heterozygous offspring genotype (0/1, 1/0, 0|1, or 1|0). At the trio level, families for which one or more members could not be linked to the current rare-disease analysis LabKey release were excluded, as were trios with fewer than 30 or more than 150 DNMs and trios without exactly one father and one mother. After filtering, 11,963 trios (11,963 probands; 6,783 male, 5,180 female) with a total of 865,392 DNMs were retained.

Each DNM was annotated with its population AF by matching to gnomAD v4.1 genome variants by chromosome, position, and allele, enabling the DNM enrichment analysis to be stratified by the AF of the existing population variant at the same position. Parent-of-origin was determined using Unfazed^24^, which phases DNMs to the paternal or maternal haplotype using informative read pairs and nearby phased variants. DNMs were classified as FATHER_ORIGIN, MOTHER_ORIGIN, or unphased, enabling separate analysis of paternally and maternally derived DNMs.

### GWAS variant data

GWAS-associated variants were obtained from the NHGRI-EBI GWAS Catalog^20^ (v1.0.2, Ensembl 113, release 2024-11-20). For each catalog association, the strongest-SNP rsID was resolved to GRCh38 coordinates and alleles via the Ensembl variation API (release 113). Each variant was annotated with its gnomAD AF by matching chromosome, position, reference allele, and alternative allele; variants without a matching gnomAD entry were excluded from the enrichment analysis (see Enrichment analysis). Variants were stored in sorted, tabix-indexed BED format per chromosome. GWAS depletion at gene conversion-compatible positions was assessed using the same contingency statistics framework as for gnomAD variants, pooled across all phenotypes and effect directions.

### Random variant generation

For the negative control analysis, a set of 10 million random single-nucleotide substitutions was generated uniformly across the genome. The per-chromosome variant count was scaled in proportion to chromosome length (target total 10^7^; genome size 3×10^9^ bp, autosomes plus X and Y). Positions were sampled uniformly within each chromosome; the reference nucleotide at each position was retrieved with twoBitToFa from the GRCh38 2bit file; positions where the reference was N were discarded; an alternative allele was selected uniformly at random from the three non-reference nucleotides. Per-chromosome BEDs were sorted, bgzipped, and tabix-indexed identically to the gnomAD variant files; the combined set was then restricted to the GIAB v3.5 all-mappable mask using bedtools intersect to produce the final VariantsToQuery.bed file (n = 8,953,591) used as input to the enrichment analyses.

### Enrichment analysis (contingency statistics)

For each combination of template length (k), AF interval, and genomic context, enrichment of variants at gene conversion-compatible positions was assessed using 2×2 contingency tables. Enrichment analyses were stratified by three genomic contexts: (1) all mappable regions (GRCh38_notinlowmappabilityall), (2) SDs only (GRCh38_notinlowmappabilitysegdups), and (3) non-SD mappable regions (GRCh38_notinlowmappabilitynosegdups), defined from the GIAB genome stratification v3.5^25^ intersected with the GIAB SD annotation. This framework was applied to four variant sets: gnomAD genome variants (stratified by AF), DNMs (stratified by the gnomAD AF of the existing population variant at the same position), GWAS Catalog variants (stratified by their gnomAD AF), and random control variants (unstratified). Per-chromosome counts were accumulated in non-overlapping 1 Mb windows; genome-wide totals are window-size-invariant because counts are summed across windows. Within each window, every mappable position was classified along two axes: (1) whether it is gene conversion-compatible at the given k, and (2) whether it harbours a variant in the given AF bin.

Consistent with the set-difference assignment (see below), the non-compatible group at each k excludes positions compatible at any longer template length, ensuring the control reflects the true non-compatible background. This produces the four cells of the contingency table: (gc_var, gc_novar, nongc_var, nongc_novar). Counts were summed across all windows and chromosomes (1–22, X, Y) to obtain genome-wide totals. A chi-squared test was applied to each genome-wide contingency table to assess whether variants occur at gene conversion-compatible positions more frequently than expected under uniform variant density.

#### Concordant variant test

At each gene conversion-compatible position harbouring a variant, the variant was classified as concordant if the observed nucleotide change matched any predicted donor allele (i.e. the alternative allele equals the central nucleotide of at least one matching k-mer pair), and discordant if the nucleotide change did not match any donor allele at any template length, including longer templates. This ensures that variants concordant at a different k value were excluded from the discordant count. The expected concordant rate was estimated from the variant rate at discordant nucleotide changes.

*Effect sizes* are reported as excess percentage: (observed − expected) / total variants × 100%. For positional enrichment this measures the fraction of variants in each frequency class occurring above the non-compatible background; for the concordance test, the analogous excess of concordant over discordant variants estimates the fraction specifically attributable to ectopic gene conversion. Values in parentheses in the enrichment tables report the percentage increase relative to expectation: (observed − expected) / expected × 100%.

#### Cumulative excess across template lengths

Because positions compatible at multiple template lengths are assigned exclusively to the longest compatible k by set difference, the conversion-compatible positions for each k are non-overlapping. The cumulative excess across all values of k was therefore computed by summing the excess percentages for each k within each AF bin: cumulative excess = Σ_k_ ExcessPercPos_k_, where ExcessPercPos_k_ = (observed_k_ − expected_k_) / total variants × 100%, and expected_k_ is the number of variants expected at the conversion-compatible positions for that k under the variant density observed at the non-compatible positions for that k. This summation is valid because the set-difference method ensures that each genomic position contributes to exactly one value of k.

#### GWAS depletion

GWAS variants were analysed using the same contingency framework, across all three mappability strata (all mappable regions, outside SDs, within SDs; Supplementary Table 21a–c). For each combination of template length and AF bin, the depletion of GWAS variants at conversion-compatible positions was computed as (VF_Conv_ − VF_NoConv_) / VF_NoConv_ × 100%, where VF_Conv_ is the per-position GWAS variant density at gene conversion-compatible positions and VF_NoConv_ the density at non-compatible positions at the same AF; negative values therefore denote depletion. This compares GWAS variant density between GC and non-GC positions within the GWAS Catalog itself.

### Multiple testing correction

For the gnomAD enrichment analysis, a total of 90 independent tests were performed per genomic context (10 template lengths × 9 AF bins), and Bonferroni correction was applied with m = 90. The same correction factor (m = 90) was used for the concordance tests (10 × 9 = 90 tests per genomic context) and for the GWAS depletion analysis (10 × 9 = 90 tests). For the DNM enrichment analysis, m = 90 was applied separately to three independent variant sets: all DNMs (ALL), paternally phased DNMs (FATHER_ORIGIN), and maternally phased DNMs (MOTHER_ORIGIN), each tested across 10 template lengths × 9 population AF bins; each set constitutes an independent analysis addressing a distinct biological question and was corrected independently. For the random variant analysis, variants were not stratified by AF, yielding 10 tests (one per template length); Bonferroni correction was applied with m = 10. For very small p-values below the limit of standard floating-point precision, log-scale p-values were computed using pchisq with log.p = TRUE and converted to base-10 exponents.

### Alignment-based homologous tract validation

As an independent validation of the variant enrichment, we used BLAST alignment to detect homologous tracts in the reference genome around variant positions. For each of 45,000 gnomAD variants (5,000 randomly sampled per AF interval) and 5,000 randomly placed control positions, we extracted the reference sequence in eight flanking windows around the variant, with half-widths of 20, 30, 40, 50, 60, 70, 80, and 90 bp on each side of the variant (total query lengths 41 to 181 bp), introduced the alternative allele at the central position, and searched the reference genome using blastn^26^ (word size 9, minimum 50% identity, dust filtering disabled, default e-value threshold of 10). We retained only alignments in which the subject sequence carried the alternative allele at the variant position, ensuring that each detected tract represents a potential donor capable of introducing the observed nucleotide change. For each alignment we defined the tract length as the genomic distance between the nearest flanking mismatches around the variant position, taken from the raw BLAST output. A variant was scored as having a detectable homologous tract when at least one BLAST alignment produced a tract of ≥ 21 bp. We set the detection threshold at ≥ 21 bp, the shortest template length at which the main k-mer analysis itself shows substantial enrichment (Supplementary Table 2): BLAST tracts of 17 or 19 bp occur in ≈ 76% of randomly placed control positions, providing insufficient discriminatory power, and the biological requirement of flanking homology around the variant precludes efficient ectopic gene conversion at these lengths when the variant lies near the tract boundary. For each AF bin, the number of gnomAD variants with a detectable tract was compared to the corresponding count from the random control set.

### Orthogonal long-read concordance check

As a technology-orthogonal control for short-read paralog mismapping, we recomputed the enrichment using PacBio HiFi joint-called variants from the Genomic Answers for Kids cohort^14^ (GA4K; n = 541 unrelated samples; DeepVariant and GLnexus; GRCh38; sites-only). Each autosomal gnomAD v4.1 genome SNV at AF ≥ 0.001 was assigned its set-difference longest-template-length class from positions.gcdiffs (k ∈ {17, 19, 21, 31, 41, 51, 61, 71, 81, 91} or non-compatible), its mappability stratum (allmapp, segdupmapp, nosegdupmapp; the same masks as the main analysis), and its AF bin. Counting was per position to mirror the contingency pipeline: a position contributes once per AF bin and is scored as confirmed when at least one of its gnomAD alleles in that bin is independently called at the matching (chromosome, position, REF, ALT) in GA4K. Variant-density enrichment used the identical definition as the main analysis, enr = (a / nPos_GC_) / (c / nPos_nonGC_), with the non-GC set at length k comprising positions not compatible at k (shorter-k classes plus non-compatible positions) and the denominators taken directly from the StatsSummary files; computed on gnomAD this reproduces Supplementary Tables 3 and 9 exactly, and the HiFi enrichment applies the same formula and denominators with GA4K SNVs as the variant source. The AF-adjusted mismatch test computed, per mappability stratum and k, the inverse-variance pooled difference in HiFi mismatch rate (1 minus the confirmation rate) between GC variants (longest_k = K) and non-GC variants (longest_k ≠ K) across AF bins, with Wilson 95% intervals on per-rate estimates and Newcombe intervals on the difference; the worst-case mismapping-driven enrichment inflation was bounded as 1 / (1 minus the upper 95% CI of this difference).

### Per-generation ectopic conversion rate

The per-generation rate of ectopic gene conversion-driven allele change (r_gc_) was estimated from the DNM contingency data. For each combination of template length k and mappability stratum, the excess of concordant DNMs at conversion-compatible positions over the discordant background was computed as:

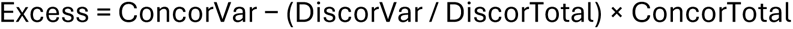

where ConcorVar and DiscorVar are the number of concordant and discordant DNMs at conversion-compatible positions (i.e. DNMs whose alternative allele matches or does not match any predicted donor allele, respectively), and ConcorTotal and DiscorTotal are the corresponding total numbers of GC-compatible positions at which concordant or discordant nucleotide changes are possible (Supplementary Tables 10 and 11, Concor. and Discord. columns). The ratio DiscorVar / DiscorTotal is the local background DNM rate from discordant changes at GC-compatible positions; using this in-region baseline rather than the rate at non-compatible positions controls for regional differences in mutation rate between conversion-compatible regions and the rest of the genome. The per-site per-generation rate was then computed as:

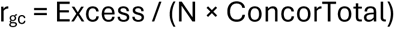

where N = 11,963 trios. As a consistency check, r_gc_ was independently estimated using the published per-nucleotide DNM rate^27^ (μ = 1.2 × 10^−8^ per bp per generation) as r_gc_ = (μ/3) × ((ConcorFreq − DiscorFreq) / DiscorFreq), where ConcorFreq = ConcorVar / ConcorTotal and DiscorFreq = DiscorVar / DiscorTotal, so that ConcorFreq − DiscorFreq is the excess concordant-DNM frequency at GC-compatible positions above the discordant baseline; this formulation eliminates the dependency on cohort size, and the two approaches agree to within a factor of 0.9–2.0 across strata, the ratio between them being exactly μ_eff_/(μ/3) in each stratum. The fold excess over the discordant background was computed as ConcorFreq / DiscorFreq, where ConcorFreq = ConcorVar / ConcorTotal. Here μ_eff_ denotes the locally estimated discordant DNM rate at conversion-compatible positions, used here in place of a nominal genome-wide point mutation rate (given for reference only as μ/3 ≈ 4 × 10^−9^ per generation). Specifically, μ_eff_ was estimated empirically for each template length and mappability stratum from the discordant DNM density at the same conversion-compatible positions, as μ_eff_ = DiscorVar / (N × DiscorTotal); the resulting values range from 3.8 × 10^−9^ to 8.1 × 10^−9^ and are reported in Supplementary Table 13, column J. Using this in-stratum empirical estimate rather than a fixed constant controls for regional differences in mutation rate between conversion-compatible regions and the rest of the genome. Accordingly, r_gc_/μ_eff_ is expressed relative to this locally estimated rate. Because DiscorTotal spans two of the three possible alternative alleles at each position whereas the concordant excess concerns a single allele, this normalisation is conservative and differs from a per-base-pair normalisation; expressing the excess relative to a single-allele rate would raise the fold excess by up to approximately two-fold. Ninety-five percent confidence intervals for the fold excess were obtained via the delta method on the log scale: log(fold) ± 1.96 × √(1/ConcorVar + 1/DiscorVar). Confidence intervals for r_gc_/μ_eff_ are derived from the fold excess CI as fold CI − 1; because r_gc_/μ_eff_ = fold excess − 1 is a linear transformation, this shift preserves the delta-method coverage exactly and is reported in Supplementary Table 13 columns O–P. Confidence intervals for r_gc_ were derived from the Poisson variance of the excess: Var(Excess) = ConcorVar + (ConcorTotal/DiscorTotal)^2^ × DiscorVar. For k = 17 within SDs, the normal approximation yields a negative r_gc_ lower bound (−3.001 × 10^−10^) because the true excess (26.4) is small relative to count noise (√ConcorVar ≈ 67.9); the exact Poisson CI on ConcorVar (lower bound = 4,476.9) likewise falls below the background count (4,582.6), confirming that clipping to 0 is appropriate.

The r_gc_/μ_eff_ lower bound at this position is also clipped to 0. Analyses were performed separately for three mappability strata (all mappable, SDs only, and non-SD mappable regions) and for gnomAD AF < 10^−5^ to restrict to truly de novo events. This rate represents the rate at which ectopic gene conversion produces a detectable allele change, not the underlying conversion initiation rate, which is substantially higher because most conversion tracts fall in homozygous regions or miss heterozygous sites.

### Non-overlapping 1-Mb window analysis

To assess the spatial distribution of gene conversion enrichment across the genome and identify outlier peaks (Figure 3, Supplementary Table 14), enrichment was computed in non-overlapping 1-Mb windows from the same 2×2 contingency table as the genome-wide analysis, with a pseudocount of 0.5 added to each cell to avoid undefined ratios. To ensure consistency with the genome-wide analysis, the non-compatible control group within each window was corrected by subtracting positions that are gene conversion-compatible at any longer template length, so that the control group contains only positions not compatible at any k value. Windows with zero gene conversion-compatible positions or zero non-compatible positions after correction were excluded. Each window was annotated with its distance to the nearest centromere and nearest telomere using GRCh38 cytoBand coordinates^28^ (UCSC acen bands), and classified as pericentromeric (within 5 Mb of the centromeric boundary), subtelomeric (within 5 Mb of the chromosome end), or interstitial. The per-chromosome distribution histograms in Supplementary Figure 3 are computed without a pseudocount, with windows lacking variants in either class excluded (see Supplementary Figure 3 caption).

#### Peak identification

For each combination of template length and AF, log_2_(OR) values across all autosomes and sex chromosomes were used directly. Outlier windows were defined as those exceeding the genome-wide mean + 2 standard deviations. Within each (k, AF) stratum independently, adjacent outlier windows on the same chromosome were merged into contiguous peak regions, yielding 529 within-stratum peak calls in total across the 90 strata. Peak regions were then collapsed across strata on each chromosome by merging overlapping and adjacent (contiguous 1 Mb) windows into 193 distinct genomic regions. Each outlier 1-Mb window was required to contain at least 10 gene conversion-compatible variants before merging. Each peak was annotated with its segmental-duplication overlap and classified into one of three categories: “segdup” (segmental-duplication coverage > 50% of the peak width), “partial” (0 < coverage < 50%), or “none” (no overlap). Each peak was further classified by chromosomal region (pericentromeric, subtelomeric, or interstitial). To confirm that the genome-wide enrichment signal is not driven by outlier regions, enrichment was recomputed after excluding the 556 individual 1-Mb outlier windows (corresponding to the 529 within-stratum peak calls), for each combination of template length and AF (Supplementary Table 15).

### Recombination hotspot analysis

To test whether enrichment at conversion-compatible positions is elevated within meiotic recombination hotspots, each genomic position was classified as inside or outside a recombination hotspot. Hotspots were defined as regions where the deCODE sex-averaged recombination rate^29^ was at least 10 cM/Mb, derived from the UCSC recombAvg bigWig track by converting to bedGraph format, thresholding at ≥ 10 cM/Mb, and merging adjacent intervals with bedtools. For each combination of template length (k ∈ {17, 19, 21, 31, 41, 51, 61, 71, 81, 91}) and AF bin, the log_2_ odds ratio for variant enrichment at conversion-compatible positions was computed separately inside and outside hotspots, restricted to autosomes. The Woolf test (z-statistic comparing log odds ratios inside versus outside, weighted by their standard errors) assessed whether enrichment differed significantly between hotspot and non-hotspot regions; Bonferroni correction was applied with m = 90 (10 template lengths × 9 AF bins). Per-template-length percentages reported in the Results (e.g. “88% stronger inside hotspots at k = 91”) are geometric-mean fold differences, computed as 2^Δlog^_2_^(OR)^ − 1, where the mean is taken across AF bins and Δlog_2_(OR) = log_2_OR_inside − log_2_OR_outside. The arithmetic ratio of mean odds ratios yields a somewhat different value at the longest templates (∼100% at k = 91).

Concordance inside and outside hotspots was computed as the proportion of variants at conversion-compatible positions whose nucleotide change matches the predicted donor allele.

### Donor-acceptor distance analysis

To assess whether conversion efficiency depends on the physical distance between donor and acceptor sequences, we extracted all donor-acceptor pairs from the gene conversion coordinate files. Pairs were filtered to 1:1 relationships (exactly one acceptor coordinate and one donor coordinate per entry) and deduplicated by canonical ordering (smaller coordinate first), as each pair appears twice in the raw data. To isolate the effect of each template length, pairs were filtered to those exclusive to each k value using the same set-difference method as the positional analysis: for each pair at template length k, both positions were required to be absent from all higher k values, ensuring that each pair contributes to exactly one template length. For each pair, the genomic distance was computed for intra-chromosomal pairs or classified as inter-chromosomal. Intra-chromosomal pairs were binned by distance: <1 kb, 1–10 kb, 10–100 kb, 100 kb–1 Mb, 1–10 Mb, 10–100 Mb, >100 Mb. Each pair was annotated with gnomAD v4.1 variant data at both the donor and acceptor positions. For each pair, the per-pair AF was defined as the maximum AF among conversion-consistent variants at the donor and acceptor positions. Mean AF per distance bin was the average of these per-pair AF values; pairs without a conversion-consistent variant at either position were excluded from the mean. Intra-versus inter-chromosomal comparisons used weighted mean AF across all intra-chromosomal distance bins. To quantify the joint predictive power of template length and distance, we fit an unweighted linear regression of mean AF on k and log_10_(distance midpoint) across all k × distance bin combinations (intra-chromosomal only) and assessed accuracy by leave-one-k-out cross-validation, reporting the cross-validated R^2^.

### GC-biased gene conversion analysis

To assess GC-biased resolution of heteroduplex intermediates, we classified all conversion-consistent variants at one-to-one donor-acceptor pairs as weak-to-strong (W→S: A/T reference to G/C alternate) or strong-to-weak (S→W: G/C reference to A/T alternate), using gnomAD v4.1 AFs. For each template length k, we computed mean AF separately for W→S and S→W variants and their ratio. To quantify the shift in composition with AF, we computed the fraction of W→S variants among all W→S + S→W variants at cumulative AF thresholds (AF ≥ 0, 0.0001, 0.001, 0.005, 0.01, 0.05, 0.1, 0.5).

### Linkage disequilibrium analysis

To test whether recurrent gene conversion disrupts LD, we compared mean r^2^ between variants at gene conversion-compatible positions (“GC variants”) and frequency-matched variants at non-compatible positions (“non-GC variants”). Phased genotypes were obtained from the 1000 Genomes high-coverage dataset^19^ (3,202 samples sequenced at 30× coverage, with pedigree-based phasing as provided by the dataset). The analysis was restricted to 1:1 paralog pairs (positions for which exactly one donor and one acceptor sequence are detectable in the genome at template length k), so that each GC focal has an unambiguous donor and the identity of the k-flanking sequence between donor and acceptor is well-defined. For each combination of mappability stratum, template length k, and AF bin, gnomAD variants at gene conversion-compatible positions were classified as GC variants and the remainder as non-GC variants. To manage computational cost, a maximum of 5,000 variants per chromosome were randomly sampled per variant class when the number exceeded this threshold.

For each chromosome, all GC and non-GC variants were extracted from the 1000 Genomes VCF, restricted to SNVs with A/C/G/T alleles (PLINK --snps-only just-acgt). PLINK v1.9^30^ was used to compute pairwise r^2^ between all variant pairs within each focal variant’s window (--r2 --ld-window-kb {window} --ld-window 99999 --ld-window-r2 0), where window was set to 10, 25, 50, or 100 kb. For each focal variant, mean r^2^ was computed across all neighbouring variants within the specified window. For GC focals, partner pairs at genomic distance ≤ (k−1)/2 bp from the focal position were excluded from the per-focal mean, because the flanking k-mer is by definition identical between donor and acceptor sequence and therefore cannot be modified by the gene conversion event itself. No such exclusion was applied to non-GC focals, which have no k-flanking constraint. For each combination of mappability stratum, population, window size, template length, and AF bin, the mean of per-variant mean r^2^ values was compared between GC and non-GC variant sets using a two-sided Wilcoxon rank-sum test. Non-GC control variants were drawn from the same AF bin as the conversion-compatible variants, with the same per-chromosome cap applied to both groups. Combinations with fewer than 30 variants in either group were excluded. Analyses were performed separately for EUR (633 samples) and ALL (3,202 samples) population groups.

#### Recombination rate stratification

To test whether the LD difference between conversion-compatible and control variants is confounded by local recombination rate, we stratified variants by recombination rate and repeated the LD comparison within each stratum. Per-variant recombination rates were obtained from the deCODE sex-averaged recombination map^29^, provided as a bedGraph file (recombAvg.bedGraph; 1,137,559 genomic intervals). Each variant was annotated with the recombination rate of its containing interval using positional overlap lookup. Variants were binned into six recombination rate categories: 0–0.5, 0.5–1, 1–2, 2–5, 5–10, and ≥10 cM/Mb. Within each recombination rate bin, the mean r^2^ of conversion-compatible variants was compared to that of frequency-matched controls using a Wilcoxon rank-sum test (minimum 10 variants per group).

#### GWAS variant depletion

To test whether conversion-compatible positions are under-represented among trait-associated common variants, we compared the rate of GWAS Catalog^20^ variants overlapping conversion-compatible positions to the rate expected from frequency-matched gnomAD variants. For each combination of template length k and AF bin, depletion (%) was computed as (VF_Conv_ − VF_NoConv_) / VF_NoConv_ × 100, where VF_Conv_ and VF_NoConv_ are GWAS variant densities at gene conversion-compatible and non-compatible positions, respectively; negative values therefore denote depletion. Significance was assessed by χ^2^ test (Supplementary Table 21). The analysis was performed across all three mappability strata: all mappable regions (main analysis, Figure 5b), outside SDs, and within SDs (Supplementary Table 21a–c).

### Computational implementation

All analyses were implemented in Perl v5.32, R v4.3.2 (with packages ggplot2, dplyr, tidyr), and bash. Genomic data were stored in sorted, bgzip-compressed, tabix-indexed BED format for efficient random-access retrieval (htslib/tabix v1.15.1^31^). BED file operations used BEDTools v2.30.0^32^. Reference sequence extraction used twoBitToFa (UCSC Genome Browser^28^ utilities). BLAST-based tract detection used blastn from NCBI BLAST+ v2.5.0^26^. LD was computed using PLINK v1.9^30^. Parent-of-origin phasing of DNMs used Unfazed v1.0.2^24^, with BCFtools v1.16^31^ used to subset the trio VCFs for Unfazed input.

## Data availability

The GRCh38 human reference genome was obtained from the UCSC Genome Browser. Data from the National Genomic Research Library (NGRL) used in this research are available within the secure Genomics England Research Environment. Access to NGRL data is restricted to adhere to consent requirements and protect participant privacy. Data used in this research include: the 100,000 Genomes Project de novo variant call set produced by the Genomics England de novo variant caller, accessed via LabKey (release V9); rare-disease programme participant and pedigree records, accessed via LabKey, used to assemble parent-offspring trios; the corresponding trio genome small-variant VCFs, used for de novo variant filtering and as input to Unfazed for parent-of-origin phasing; and the per-sample read alignments (BAM/CRAM) for the trio members, used by Unfazed for read-backed phasing. Derived summary statistics underlying the de novo mutation analyses (the genome-wide and per-stratum contingency counts, concordance statistics, and per-generation rate estimates) were generated within the Research Environment and exported via the Genomics England Airlock process. Access to NGRL data is provided to approved researchers who are members of the Genomics England Research Network, subject to institutional access agreements and research project approval under participant-led governance. For more information on data access, visit: genomicsengland.co.uk/research

Genome stratification files and SD annotations were obtained from GIAB v3.5. Population variant data were obtained from gnomAD v4.1. Phased genotypes were obtained from the 1000 Genomes high-coverage dataset. GWAS-associated variants were obtained from the NHGRI-EBI GWAS Catalog (v1.0.2, Ensembl 113). Recombination rates were obtained from deCODE genetics via the UCSC Genome Browser. Centromere coordinates were obtained from the UCSC cytoBand track. PacBio HiFi joint-called variants used for the orthogonal long-read validation were obtained from the Genomic Answers for Kids (GA4K) cohort as a sites-only release; access to GA4K data is governed by the study authors and described in the cited reference.

## Code availability

All custom code used in this study is publicly available at github.com/WouterSteyaert/EctopicGeneConv. The repository contains the complete analysis pipeline, from the identification of gene conversion-compatible positions to the enrichment, DNM, per-generation conversion-rate, and LD analyses.

## Acknowledgements

We gratefully acknowledge the participants of the National Genomic Research Library (NGRL), whose contributions made this research possible. Secure access to the NGRL under project ID 1017 was provided by Genomics England, which delivers the NGRL in partnership with NHS England, and is wholly owned by the UK Department of Health and Social Care. The NGRL contains participants’ health data collected by the NHS as part of their care, along with samples and data from their participation in research, for which fully informed consent has been obtained. This includes genomic and clinical data provided through the NHS Genomic Medicine Service, as well as data obtained through research studies, including the 100,000 Genomes Project and the Generation Study, both of which are delivered in partnership with the NHS, and from other research cohorts involving external collaborators.

The computational resources (Stevin Supercomputer Infrastructure) and services used in this work were provided by the VSC (Flemish Supercomputer Center), funded by Ghent University, FWO and the Flemish Government (department EWI).

## Author contributions

W.S. conceived the study, developed the methodology, performed all analyses, and wrote the manuscript.

## Competing interests

The author declares no competing interests.

## Ethics approval

This study analysed DNM data from the 100,000 Genomes Project, held within the National Genomic Research Library (NGRL). The 100,000 Genomes Project has standing research ethics approval from the East of England – Cambridge South Research Ethics Committee (REC reference 14/EE/1112); all participants provided written informed consent for their data to be used in research, with secondary analyses covered under this approval. Access for the present study was approved by Genomics England under NGRL Research Registry project ID 1017.

## Supplementary Figure Legends

**Supplementary Figure 1: Variant-density enrichment reproduces in PacBio HiFi long reads.** Variant-density enrichment in gnomAD (short-read) and GA4K HiFi long-read calls per mappability stratum, template length k, and allele-frequency bin, log scale. Both sources use the identical position partition and the same per-(stratum, k, AF) position denominators (nPos_GC_, nPos_nonGC_) as the main gnomAD enrichment analysis (Methods). The plot shows the five AF bins where both technologies are cohort-saturated (AF ≥ 0.005); the lowest AF bin (0.001–0.005) is omitted from the visualization because cohort detection at this AF saturates at ∼66% in the 541 GA4K samples, but it is included as the sixth stratum in the inflation pooling shown in Supplementary Figure 2 (six AF strata, AF 0.001–1). The HiFi line tracks the gnomAD line throughout, and within segmental duplications at long k the HiFi enrichment lies at or above the gnomAD enrichment, opposite to the direction expected if short-read mismapping had inflated the gnomAD signal.

**Supplementary Figure 2: Worst-case enrichment inflation from short-read mismapping.** Per (mappability stratum, template length k), the worst-case enrichment-inflation factor derived from the AF-adjusted Δ mismatch (HiFi mismatch rate of GC minus non-GC variants), inverse-variance pooled across six allele-frequency strata (AF 0.001–1) with Wilson 95% intervals on per-rate estimates and Newcombe intervals on the difference; the plotted factor is 1 / (1 − upper 95% CI of Δ). The factor is at most 1.009× (k = 21, within SDs), negligible against the up-to-14-fold enrichment at long templates in segmental duplications; within those regions at k ≥ 51 Δ is negative, indicating GC variants validate at slightly higher rates than non-GC.

**Supplementary Figure 3: Distribution of enrichment across non-overlapping 1-Mb windows per chromosome.** For each chromosome, histograms show the distribution of log_2_ odds ratios across non-overlapping 1-Mb windows, faceted by four representative template lengths (k = 21, 41, 61, 91; rows) and nine allele frequency bins (columns). The grey dashed vertical line marks log_2_(OR) = 0 (no enrichment); the red vertical line marks the per-panel median. The right-most column (“> 0.5”) aggregates all variants with allele frequency above 0.5. Empty facets indicate combinations with no qualifying windows on that chromosome (e.g. extreme AF bins on chromosome Y). For this visualization, the per-window log_2_ odds ratio was computed without a pseudocount, and windows lacking variants in either class (and therefore with an undefined or infinite odds ratio) were excluded, so that the histogram reflects only windows carrying an informative enrichment signal.

**Supplementary Figure 4: GC-biased gene conversion.** (a) Mean allele frequency of W→S (weak-to-strong) and S→W (strong-to-weak) conversion-consistent variants at one-to-one donor-acceptor pairs, as a function of template length. (b) Ratio of mean allele frequency W→S / S→W as a function of template length. A ratio above 1 indicates GC-biased resolution of heteroduplex intermediates.

## Supplementary Table Legends

**Supplementary Table 1:** Number of gene conversion-compatible positions per template length. For each minimal k-mer length (k), the total number of positions in the mappable genome (GRCh38, GIAB v3.5 high-mappability regions; 2.69 Gb) where at least one paralogous k-mer pair exists that differs only at the central nucleotide. Percentages indicate the fraction of the mappable genome that is gene conversion-compatible at each k. Note that these counts are not mutually exclusive: a position compatible at k = 91 is also compatible at all shorter k values. In all downstream analyses, positions are assigned exclusively to the longest compatible k by set difference to avoid double counting (Methods).

**Supplementary Table 2:** Enrichment and concordance of genetic variants at gene conversion-compatible positions (all mappable regions). Values are shown for each combination of template length k and gnomAD v4.1 genome-wide allele frequency bin (columns). (a) Relative excess of variant density at gene conversion-compatible positions compared to non-gene conversion positions. (b) Positional excess expressed as a percentage of all variants in the genome. (c) Concordance excess. The Σ row gives the sum across all k values. All 90 individual enrichment tests are significant after Bonferroni correction (p_χ_^2^ < 2.04 × 10^−18^).

**Supplementary Table 3:** Contingency statistics and Bonferroni-corrected p-values for enrichment and concordance tests (all mappable regions). For each combination of k-mer length and allele frequency bin, the four cells of the 2×2 contingency table are shown for the positional enrichment test and the concordance test. Bonferroni-corrected p-values are reported as log_10_(p) with correction factor m = 90.

**Supplementary Table 4:** Alignment-based validation of variant enrichment within BLAST-detected homologous tracts. For each AF bin, the number of gnomAD variants (5,000 sampled per bin) and random control positions (5,000 total) with a BLAST-detected homologous tract of at least 21 bp (raw asymmetric tract length) on the reference genome with the alternative allele at the central position. The k = 17 and k = 19 thresholds were not used because short asymmetric tracts (≤ 19 bp) are abundant in random sequence and do not discriminate gnomAD variants from random controls (≈ 76% detection rate at ≥ 17 bp in both groups).

**Supplementary Table 5:** Random variant analysis (negative control). Contingency statistics and Bonferroni-corrected p-values for 8,953,591 randomly placed single-nucleotide substitutions (10 million sampled, retained after N-filtering and GIAB v3.5 mappable mask intersection; Methods) tested against gene conversion-compatible positions, stratified by mappability: (a) all mappable regions; (b) within segmental duplications. Bonferroni correction was applied separately within each stratum (m = 10 per stratum). No significant enrichment or concordance is observed at any k-mer length in either stratum.

**Supplementary Table 6:** Variant-density enrichment in gnomAD versus GA4K HiFi long-read calls per mappability stratum, template length k, and allele-frequency bin, with the HiFi/gnomAD enrichment ratio. Both sources use the identical position partition and per-cell denominators as the main contingency analysis (Methods); computed on gnomAD this reproduces the published enrichment exactly. File: v2_enrichment_compare.tsv.

**Supplementary Table 7:** HiFi confirmation rates of GC and non-GC variants per mappability stratum and template length k, pooled over the reliable allele-frequency zone (AF ≥ 0.01), with Wilson 95% confidence intervals. Confirmation is 94 to 99% in every cell; within segmental duplications at k = 21 to 41 GC validates about 1 to 2% below non-GC, consistent with lower callability of short conversion units, while at k ≥ 51 GC validates equally or better. File: v2_conf_rates_reliableAF.tsv.

**Supplementary Table 8:** AF-adjusted Δ mismatch (HiFi mismatch rate of GC minus non-GC variants) and worst-case enrichment-inflation bound per mappability stratum and template length k, inverse-variance pooled across six allele-frequency strata (AF 0.001 to 1); max inflation = 1 / (1 − upper 95% CI of Δ). The bound is at most 1.009× (k = 21, within SDs); within segmental duplications at k ≥ 51 Δ is negative, indicating GC variants validate at slightly higher rates than non-GC. File: v2_mh_pooled.tsv.

**Supplementary Table 9:** Enrichment and concordance within and outside SDs. Contingency statistics and Bonferroni-corrected p-values (m = 90) for the positional enrichment and concordance tests, stratified by genomic context (within SDs, outside SDs); the all-mappable analysis is in Supplementary Tables 2 and 3.

**Supplementary Table 10:** De novo mutation enrichment at gene conversion-compatible positions. Contingency statistics and Bonferroni-corrected p-values (m = 90) for 865,392 DNMs from 11,963 parent-offspring trios, stratified by k-mer length and gnomAD allele frequency bin of the corresponding population variant.

**Supplementary Table 11:** De novo mutation enrichment outside SDs. Same analysis as Supplementary Table 10, restricted to regions outside annotated SDs.

**Supplementary Table 12:** De novo mutation enrichment by parent of origin (autosomes only). Same analysis as Supplementary Table 10, stratified by parental origin of DNMs and restricted to autosomes to avoid confounding by hemizygous chrX inheritance in males. (a) Paternally derived. (b) Maternally derived.

**Supplementary Table 13:** Per-generation ectopic gene conversion rate. For each template length, the excess of concordant over discordant DNMs yields r_gc_, the per-site per-generation rate of conversion-driven allele change (Methods). Ratios are expressed relative to μ_eff_, the discordant DNM rate estimated locally within each stratum as DiscorVar / (N × DiscorTotal). ConcorTotal and DiscorTotal give the number of conversion-compatible positions at which the concordant and the discordant nucleotide changes, respectively, are possible. Fold excess is computed as the concordant DNM rate divided by the discordant DNM rate, where each rate is expressed per GC-compatible position (i.e. ConcorVar/ConcorTotal divided by DiscorVar/DiscorTotal); r_gc_/μ_eff_ = fold excess − 1. 95% confidence intervals for r_gc_ are based on Poisson approximation on the Excess; 95% confidence intervals for fold excess are derived via the delta method on the log scale (columns L–N). 95% confidence intervals for r_gc_/μ_eff_ are obtained as fold excess CI − 1 (columns O–P); this linear shift preserves the delta-method coverage exactly. Where the normal approximation yields a negative lower bound for r_gc_ (k = 17, within SDs: −3.001 × 10^−10^; confirmed negative also under exact Poisson CI), the lower bound is clipped to 0 (marked †; see table footnote). The corresponding r_gc_/μ_eff_ lower bound is likewise clipped to 0. (a) All mappable regions. (b) Within SDs. (c) Outside SDs.

**Supplementary Table 14:** Non-overlapping 1-Mb window enrichment peaks at gene conversion-compatible positions. (a) **529 within-stratum peak calls.** Outlier 1-Mb windows (log_2_ odds ratio exceeding the genome-wide mean by more than two standard deviations) for each combination of template length and allele frequency bin, with chromosomal location, segmental duplication overlap, and regional classification (pericentromeric, subtelomeric, interstitial). Telomere distance is computed as the distance from the window midpoint to the nearest chromosome end; negative values indicate windows whose midpoint extends marginally beyond the annotated chromosome boundary and should be interpreted as 0 Mb. (b) **193 distinct genomic regions.** Interval-merging the 529 within-stratum peak calls in (a) across all (k, AF) strata (Methods) yields 193 distinct genomic regions. For each merged region: chromosomal coordinates, the number of (k, AF) strata in which it was called, the maximum log_2_(OR) across those strata, segmental-duplication overlap, and regional classification. The percentages reported in the Results (interstitial (64%), pericentromeric (26%), subtelomeric (10%); 83% overlap annotated SDs) are computed from this sheet.

**Supplementary Table 15:** Enrichment at gene conversion-compatible positions after excluding outlier windows. For each combination of template length (k) and allele frequency bin, enrichment statistics are shown for all 1-Mb windows and after excluding the 556 individual 1-Mb outlier windows (corresponding to the 529 within-stratum peak calls in Supplementary Table 14). Mean log_2_OR: mean log_2_ odds ratio across windows. % windows positive: percentage of windows with log_2_OR > 0.

**Supplementary Table 16:** Enrichment at gene conversion-compatible positions within and outside recombination hotspots. For each combination of template length and allele frequency bin, the log_2_ odds ratio for variant enrichment at conversion-compatible positions is shown separately for positions inside and outside meiotic recombination hotspots (deCODE sex-averaged rate ≥ 10 cM/Mb), with Woolf test statistics for heterogeneity.

**Supplementary Table 17:** Allele frequency of conversion-consistent variants as a function of donor-acceptor distance. For each template length, unambiguous one-to-one donor-acceptor pairs were stratified by genomic distance into seven intra-chromosomal bins (<1 kb, 1–10 kb, 10–100 kb, 100 kb–1 Mb, 1–10 Mb, 10–100 Mb, >100 Mb) and one inter-chromosomal category. Mean allele frequency of conversion-consistent gnomAD v4.1 variants is reported per bin.

**Supplementary Table 18:** GC-biased gene conversion analysis. For each template length k, the fraction of W→S variants among all W→S and S→W variants at cumulative allele frequency thresholds. Mean allele frequencies per direction (W→S and S→W) are shown in Supplementary Figure 4.

**Supplementary Table 19:** Linkage disequilibrium at gene conversion-compatible positions. Mean r^2^ for variants at gene conversion-compatible positions (“GC variants”) and frequency-matched non-compatible positions (“non-GC variants”), for each combination of mappability stratum, population (EUR, ALL), window size (10, 25, 50, 100 kb), k-mer length, and minor allele frequency bin. Analysis restricted to 1:1 paralog pairs (positions with exactly one donor and one acceptor at template length k) to ensure unambiguous donor-acceptor identity in the k-flanking region. For each GC focal, partner variants within ±(k−1)/2 bp were excluded because the k-flanking sequence is identical between donor and acceptor by construction; PLINK was run with --snps-only just-acgt to exclude indels and multi-nucleotide variants (Methods). P-values from two-sided Wilcoxon rank-sum test; combinations with n < 30 variants per group excluded.

**Supplementary Table 20:** LD stratified by local recombination rate. Mean r^2^ for variants at GC-compatible positions versus frequency-matched non-GC controls, for each combination of mappability stratum, population (EUR, ALL), window size (10, 25, 50, 100 kb), k-mer length, allele frequency bin, and local recombination rate bin (0–0.5, 0.5–1, 1–2, 2–5, 5–10, ≥10 cM/Mb). Analysis restricted to 1:1 paralog pairs with ±(k−1)/2 bp partner exclusion and PLINK --snps-only just-acgt (Methods). Recombination rates from the deCODE sex-averaged genetic map. P-values from two-sided Wilcoxon rank-sum test; combinations with n < 10 variants per group excluded.

**Supplementary Table 21:** GWAS variant depletion at gene conversion-compatible positions. Three sheets stratified by mappability: (21a) all mappable regions; (21b) outside segmental duplications; (21c) within segmental duplications. For each combination of k-mer length and allele frequency bin, contingency statistics compare GWAS Catalog variant density at GC-compatible positions versus non-compatible positions. Depletion (under-representation) is indicated by a lower variant density at conversion-compatible positions (GC frequency < non-GC frequency); the χ^2^ test p-values, reported as Bonferroni-corrected log_10_ p-values (m = 90), give the significance of the difference; concordance statistics compare allele-direction agreement between GWAS variants and the predicted donor-to-acceptor change.

## Notes

https://github.com/WouterSteyaert/EctopicGeneConv

### Competing Interest Statement

The authors have declared no competing interest.

