## Supplementary Figures 1-4 for "Sequence architecture shapes human allele frequencies through ectopic gene conversion"

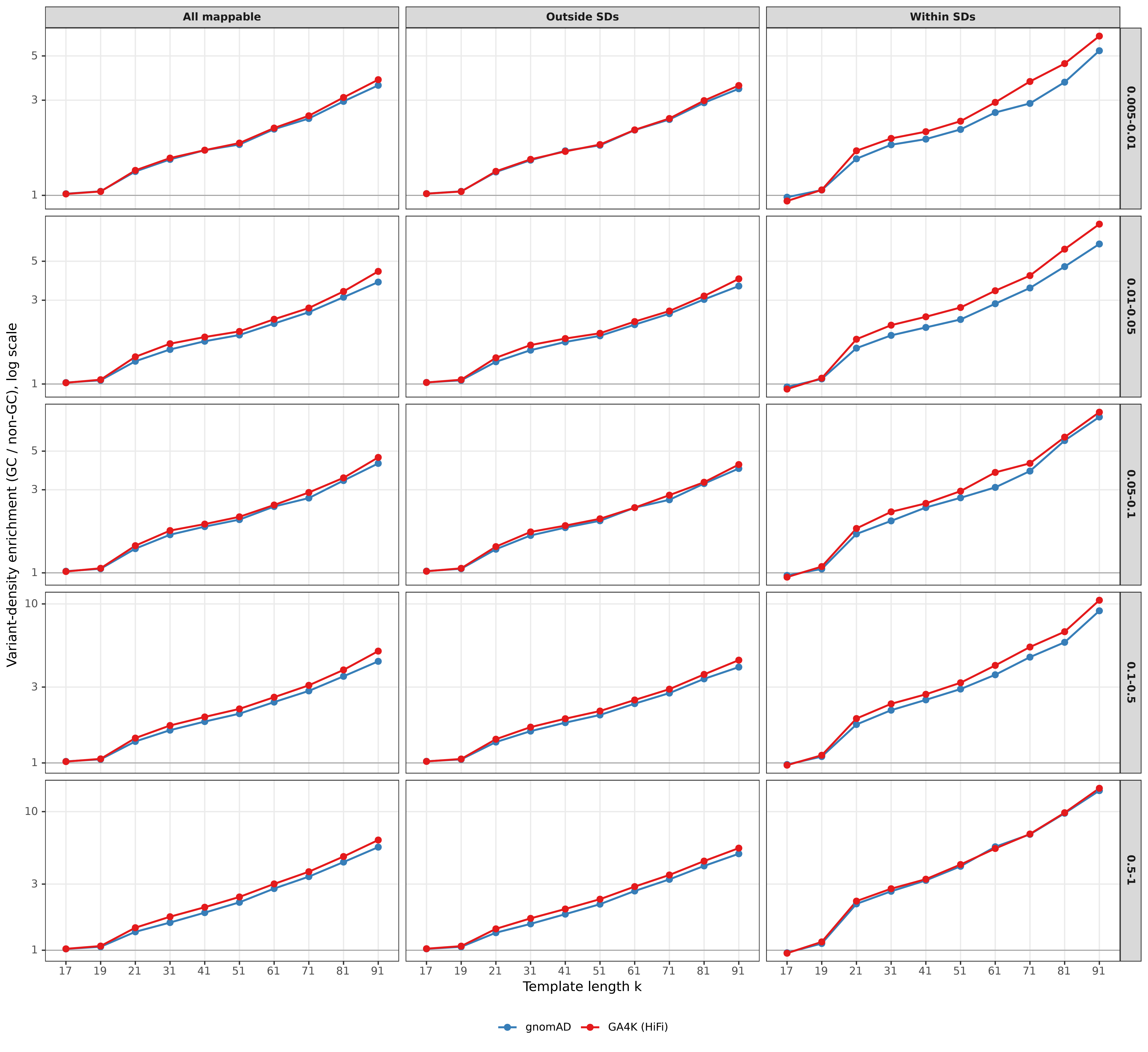

All mappable
Outside SDs
Within SDs
0.005-0.01
0.01-0.05
0.05-0.1
0.1-0.5
0.5-1
Variant-density enrichment (GC / non-GC), log scale
Template length k
17
19
21
31
41
51
61
71
81
91
1
3
5
10
gnomAD
GA4K (HiFi)

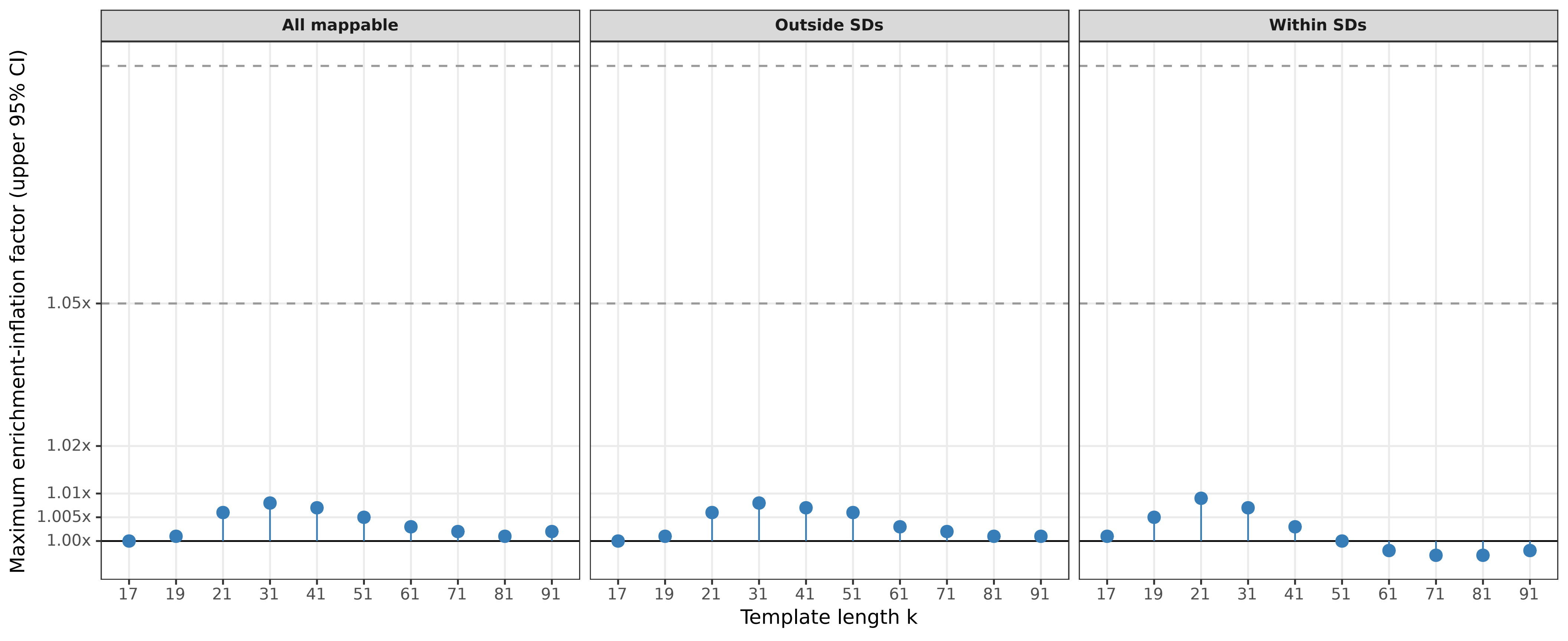
All mappable
Outside SDs
Within SDs
Maximum enrichment-inflation factor (upper 95% CI)
1.05x
1.02x
1.01x
1.005x
1.00x
17
19
21
31
41
51
61
71
81
91
Template length k

#### Chromosome 1

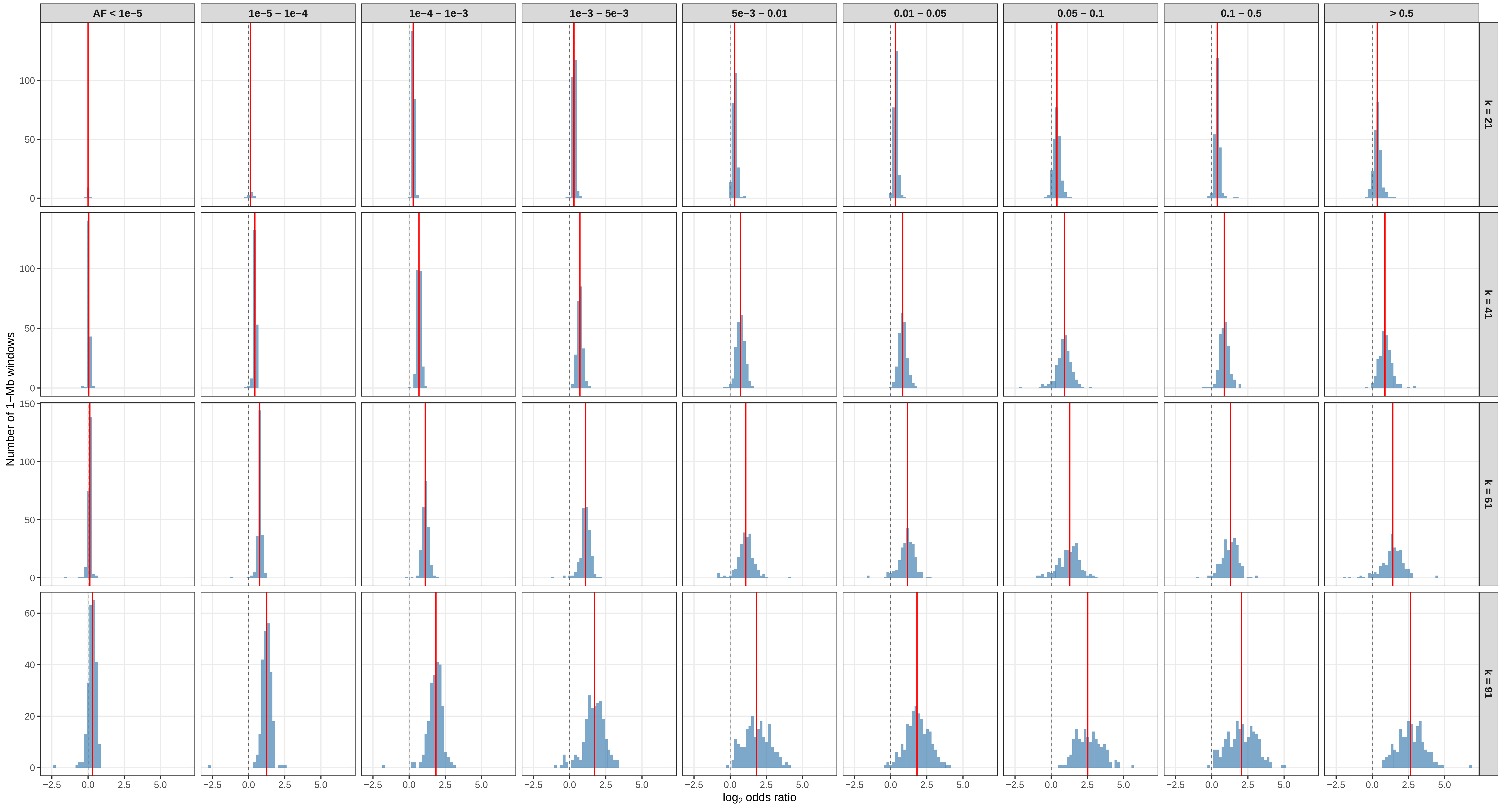

### Chromosome 2

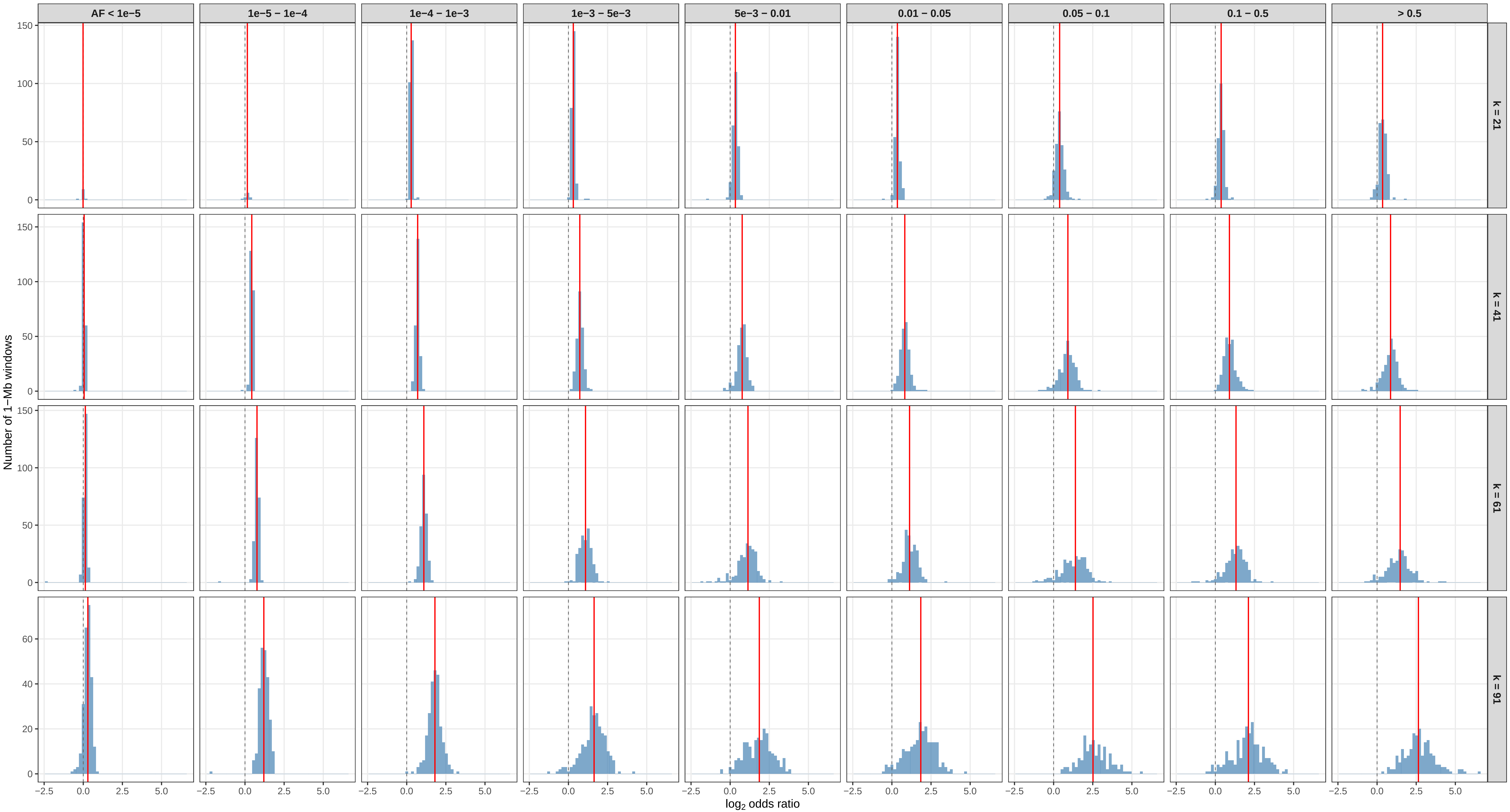

#### Chromosome 3

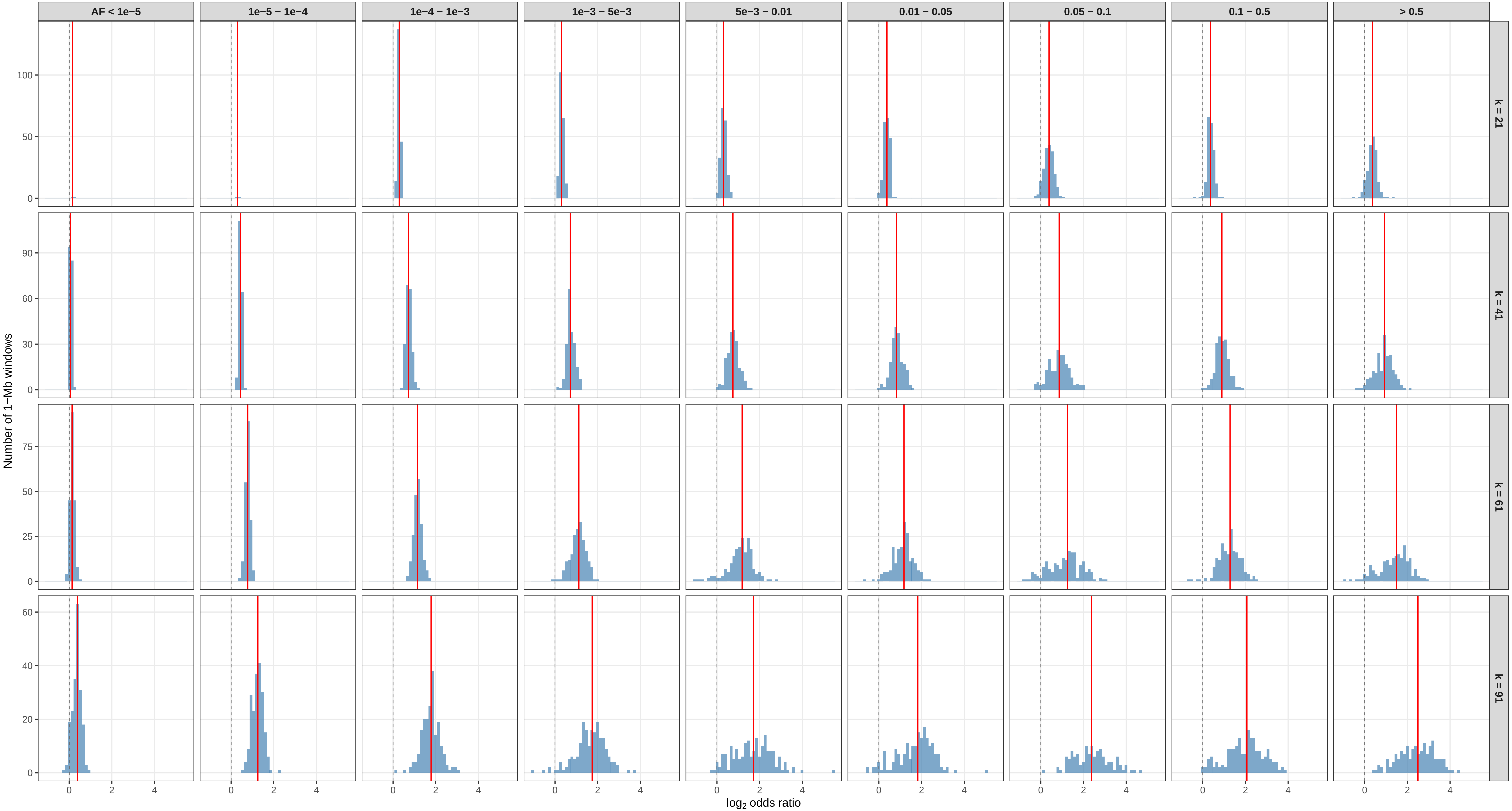

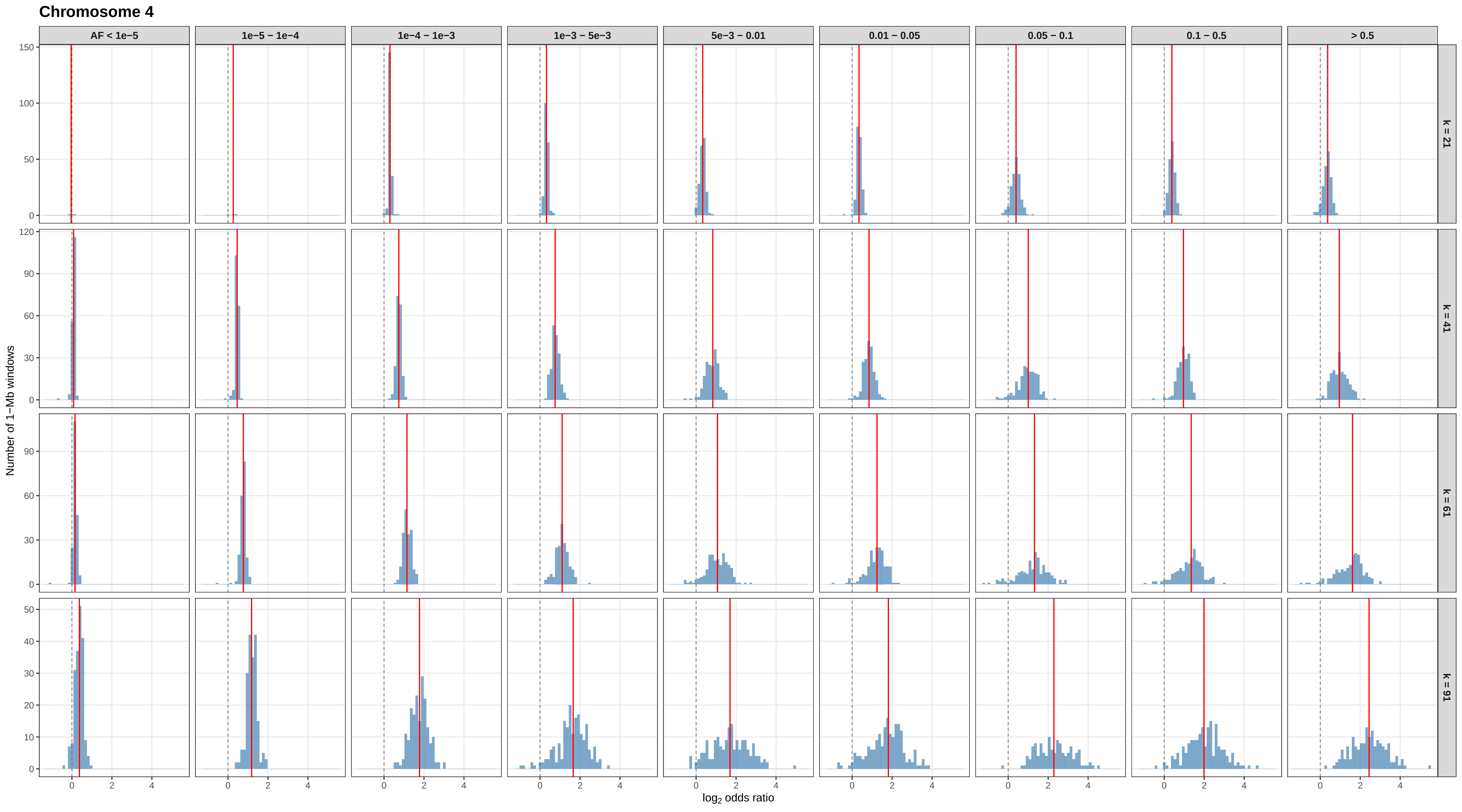

Chromosome 4
AF < 1e−5
1e−5 – 1e−4
1e−4 – 1e−3
1e−3 – 5e−3
5e−3 – 0.01
0.01 – 0.05
0.05 – 0.1
0.1 – 0.5
> 0.5
k = 21
k = 41
k = 61
k = 91
Number of 1−Mb windows
log2 odds ratio

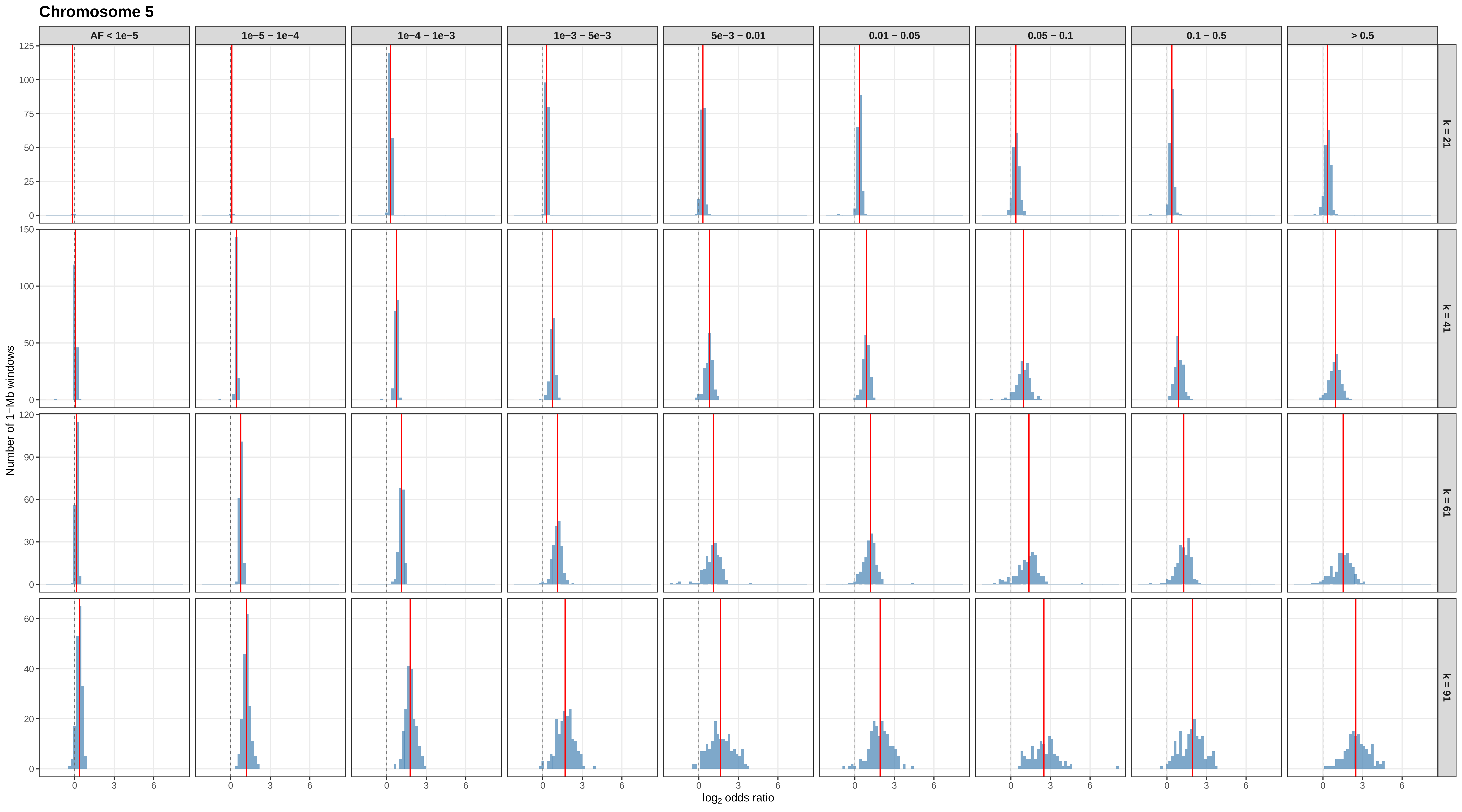

Chromosome 5
AF < 1e−5
1e−5 − 1e−4
1e−4 − 1e−3
1e−3 − 5e−3
5e−3 − 0.01
0.01 − 0.05
0.05 − 0.1
0.1 − 0.5
> 0.5
k = 21
k = 41
k = 61
k = 91
Number of 1−Mb windows
log2 odds ratio

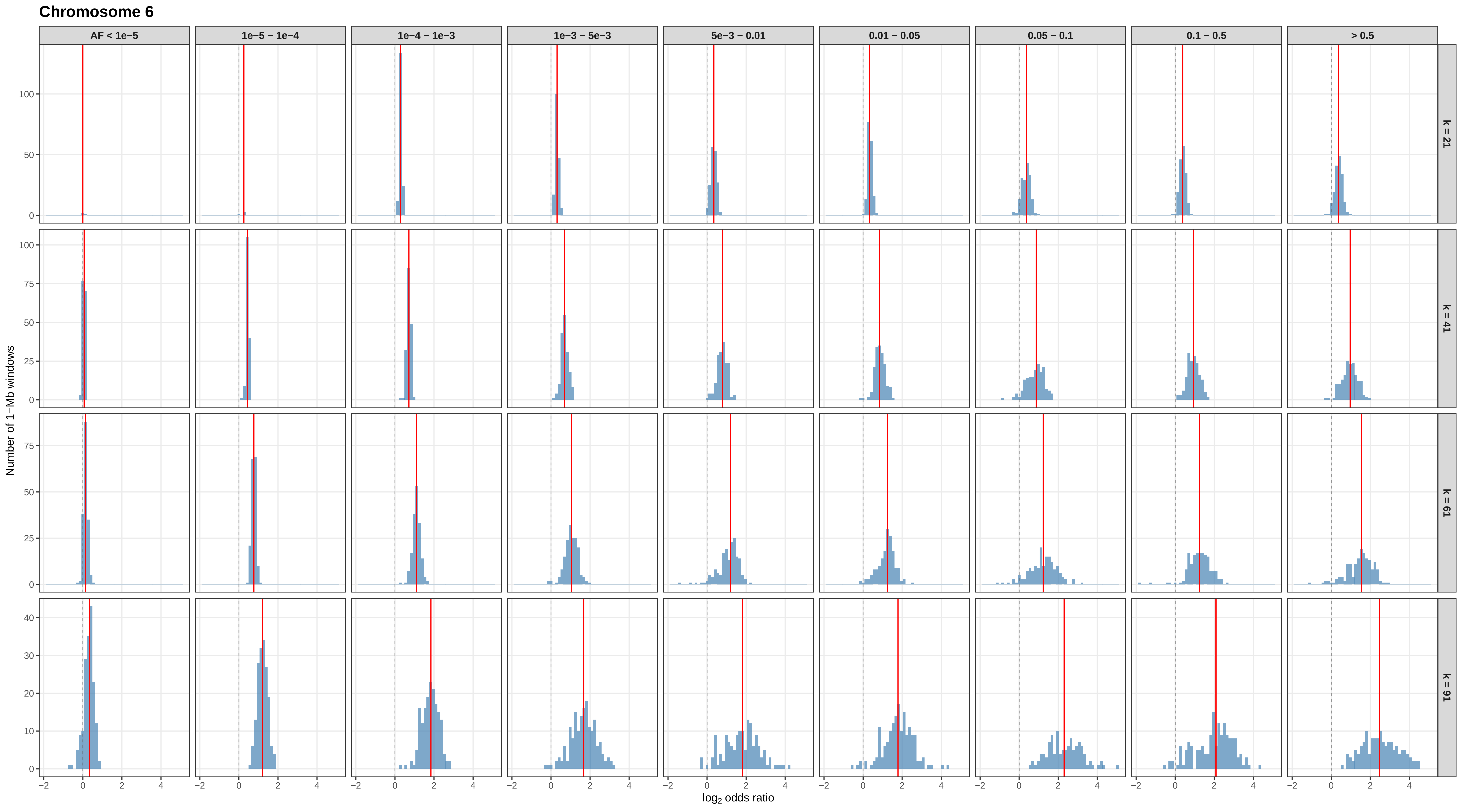

Chromosome 6
AF < 1e−5
1e−5 – 1e−4
1e−4 – 1e−3
1e−3 – 5e−3
5e−3 – 0.01
0.01 – 0.05
0.05 – 0.1
0.1 – 0.5
> 0.5
k = 21
k = 41
k = 61
k = 91
Number of 1−Mb windows
log2 odds ratio

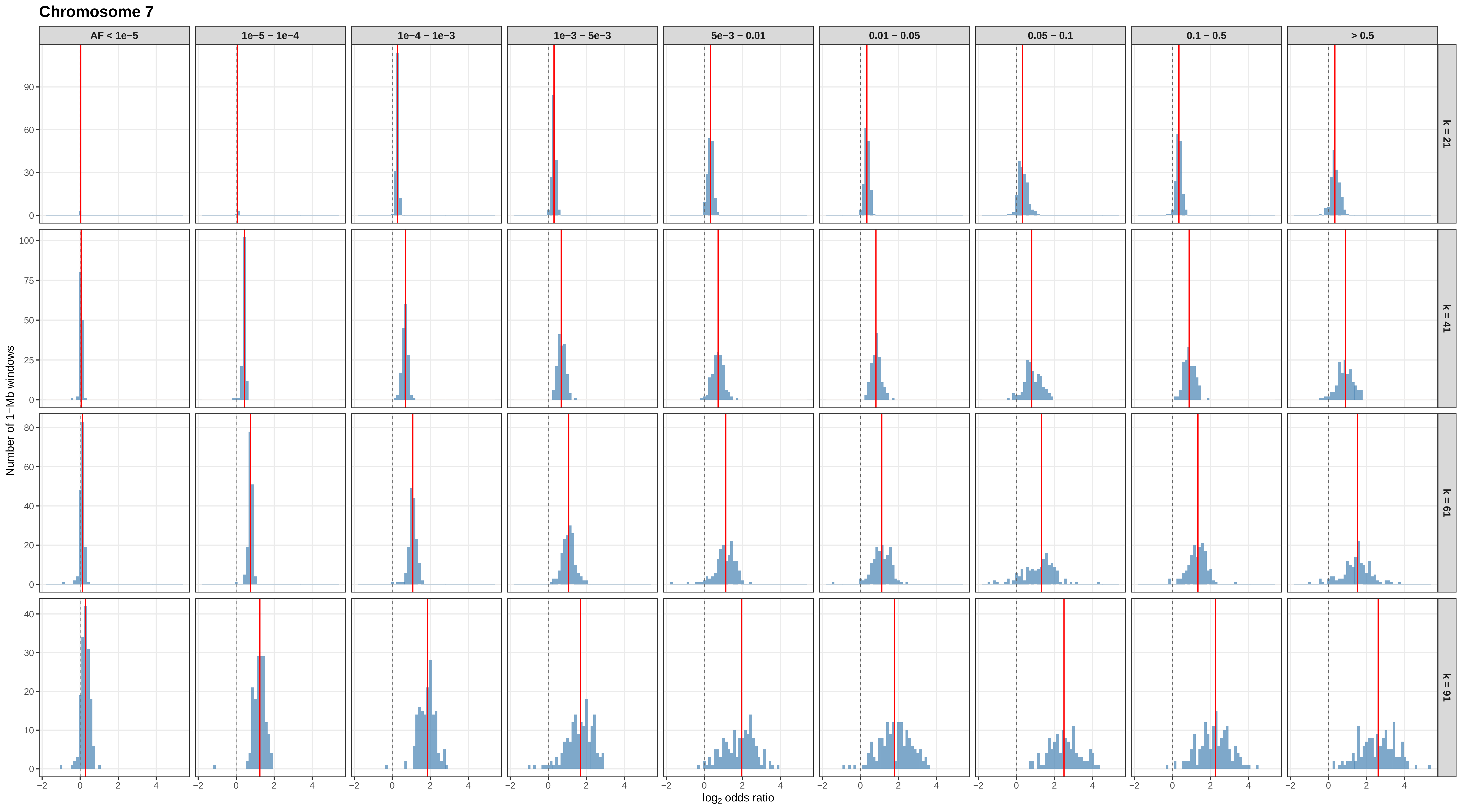
Chromosome 7
AF < 1e−5
1e−5 – 1e−4
1e−4 – 1e−3
1e−3 – 5e−3
5e−3 – 0.01
0.01 – 0.05
0.05 – 0.1
0.1 – 0.5
> 0.5
k = 21
k = 41
k = 61
k = 91
Number of 1−Mb windows
log2 odds ratio

Chromosome 8

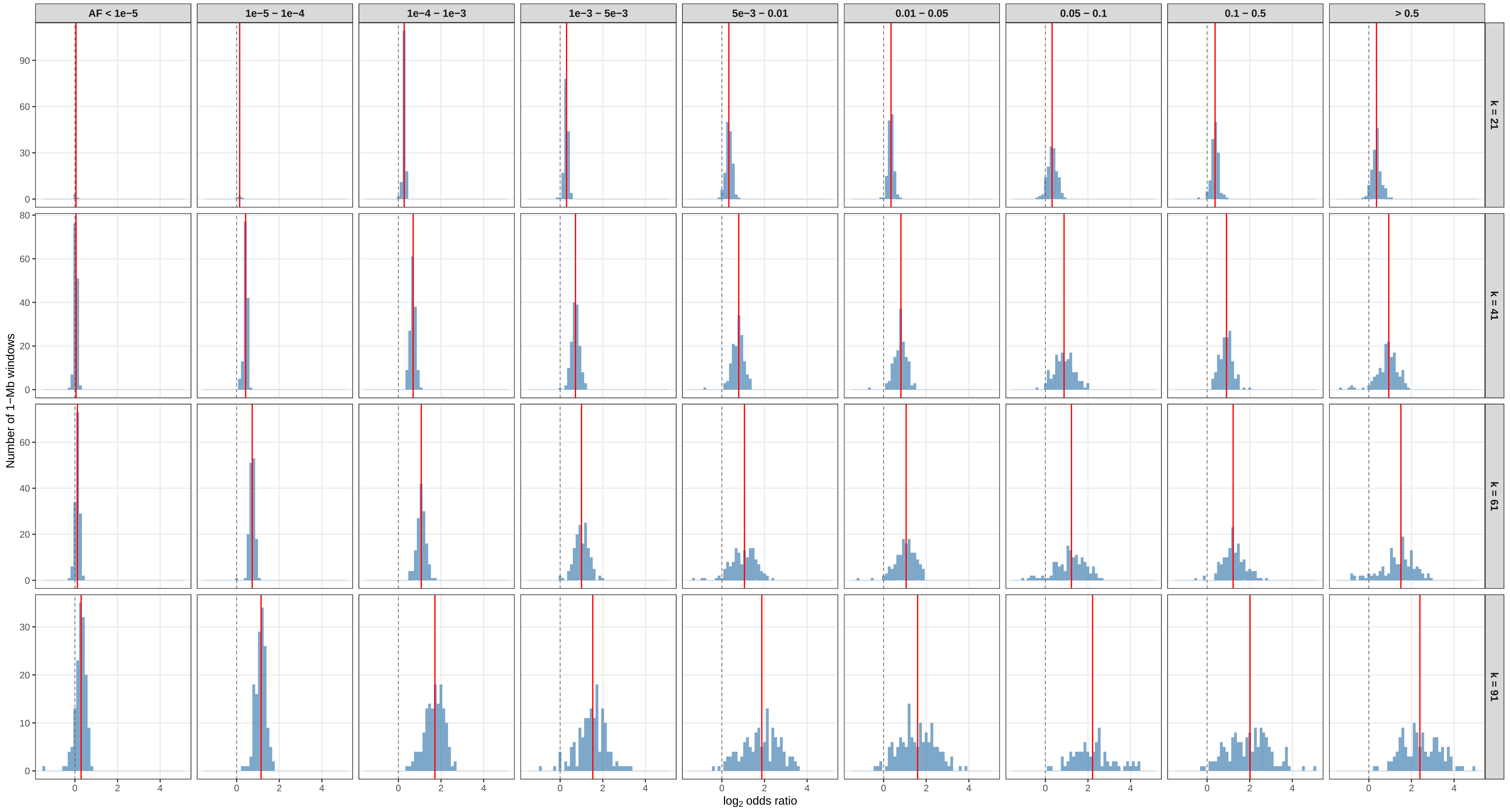

#### Chromosome 9

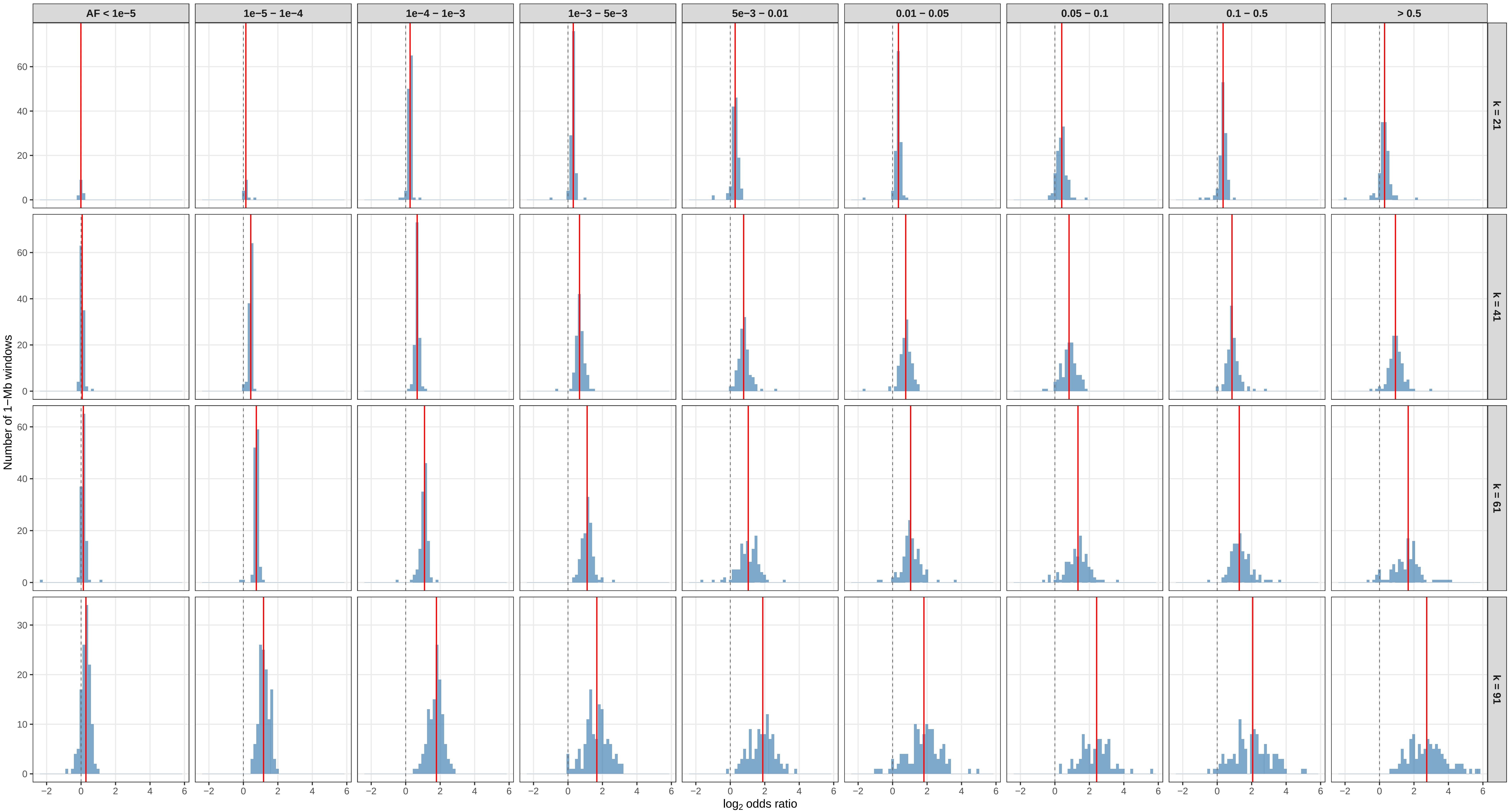

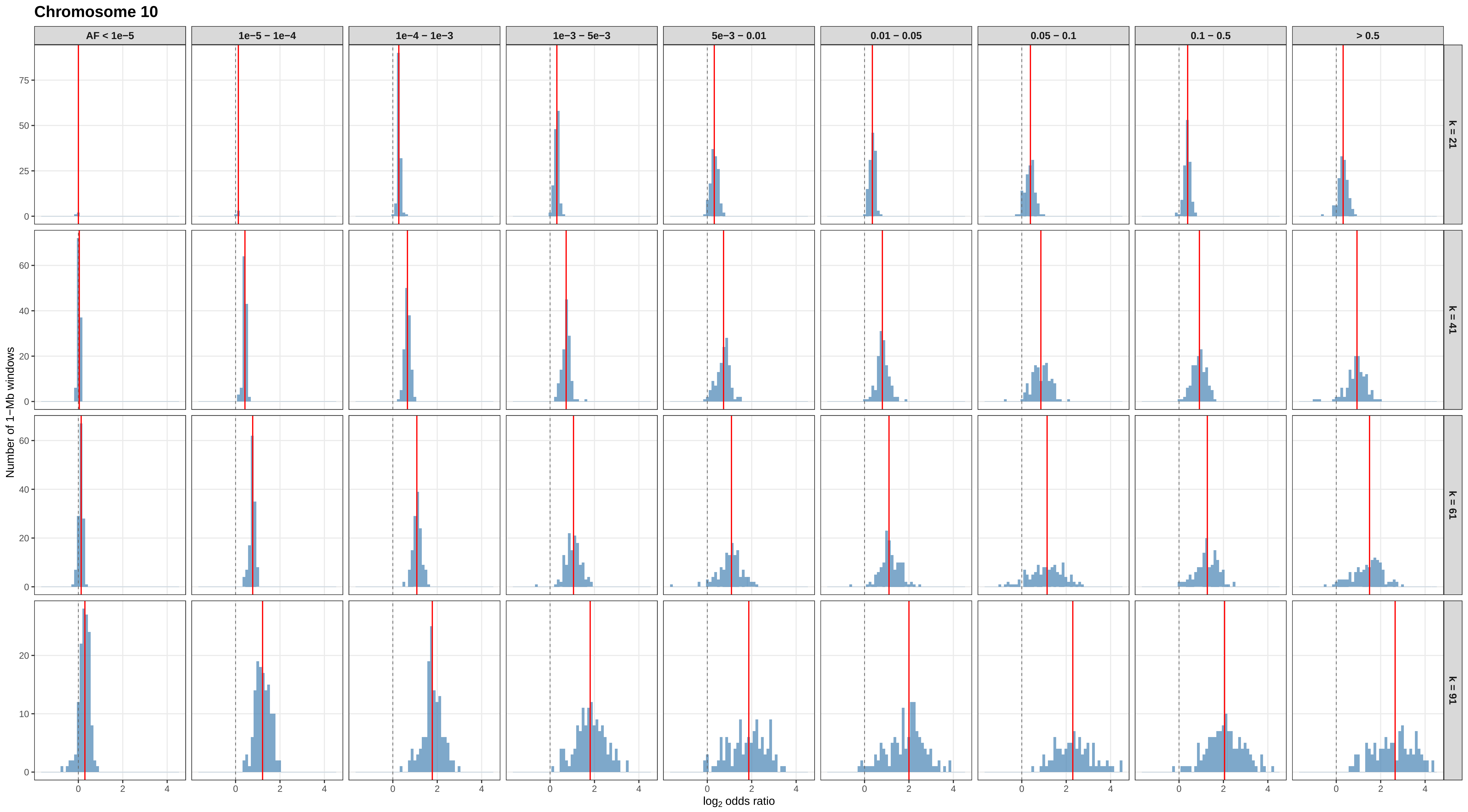
Chromosome 10
AF < 1e−5
1e−5 – 1e−4
1e−4 – 1e−3
1e−3 – 5e−3
5e−3 – 0.01
0.01 – 0.05
0.05 – 0.1
0.1 – 0.5
> 0.5
k = 21
k = 41
k = 61
k = 91
Number of 1−Mb windows
log2 odds ratio

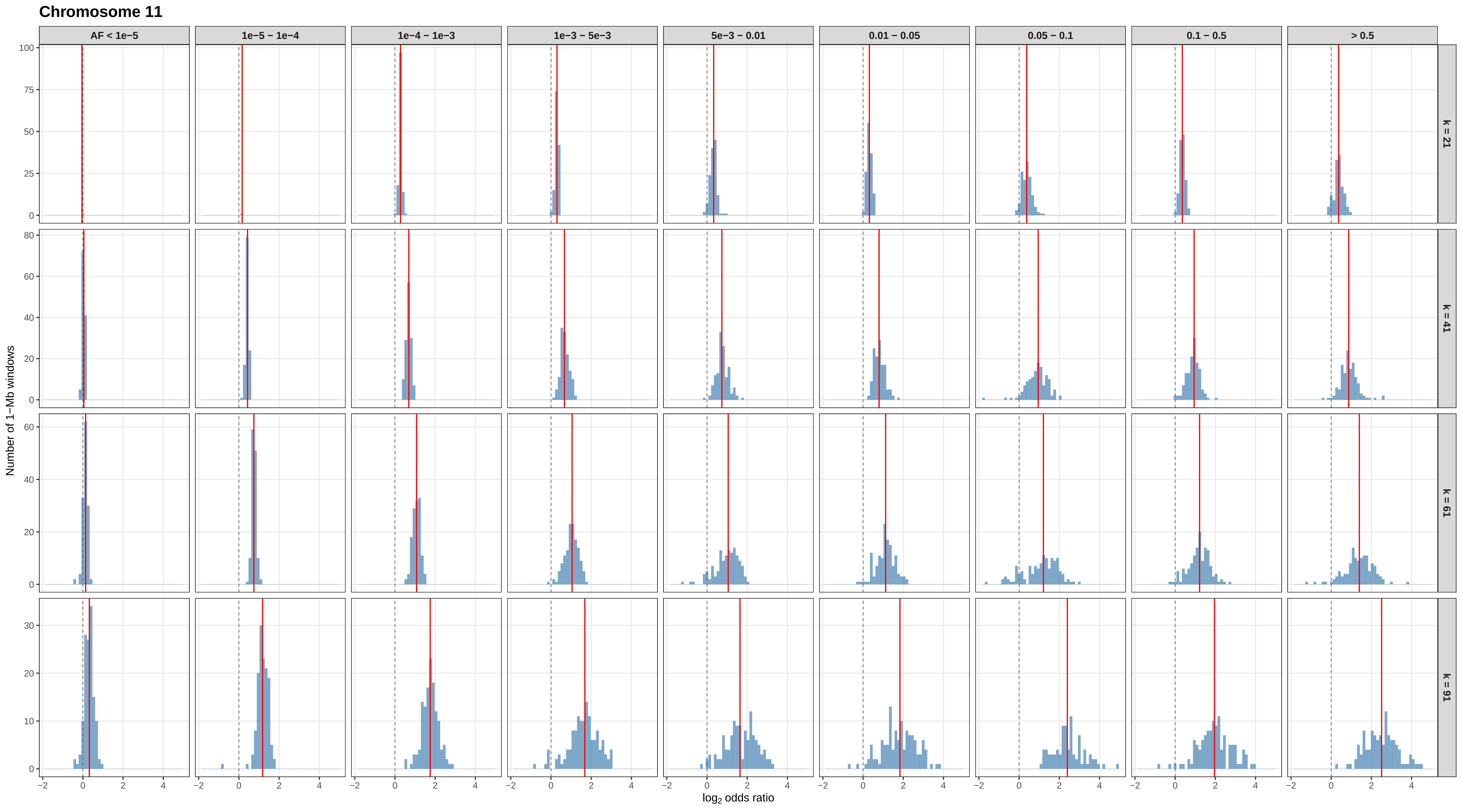
Chromosome 11
AF < 1e−5
1e−5 − 1e−4
1e−4 − 1e−3
1e−3 − 5e−3
5e−3 − 0.01
0.01 − 0.05
0.05 − 0.1
0.1 − 0.5
> 0.5
k = 21
k = 41
k = 61
k = 91
Number of 1−Mb windows
$\log_2$ odds ratio

#### Chromosome 12

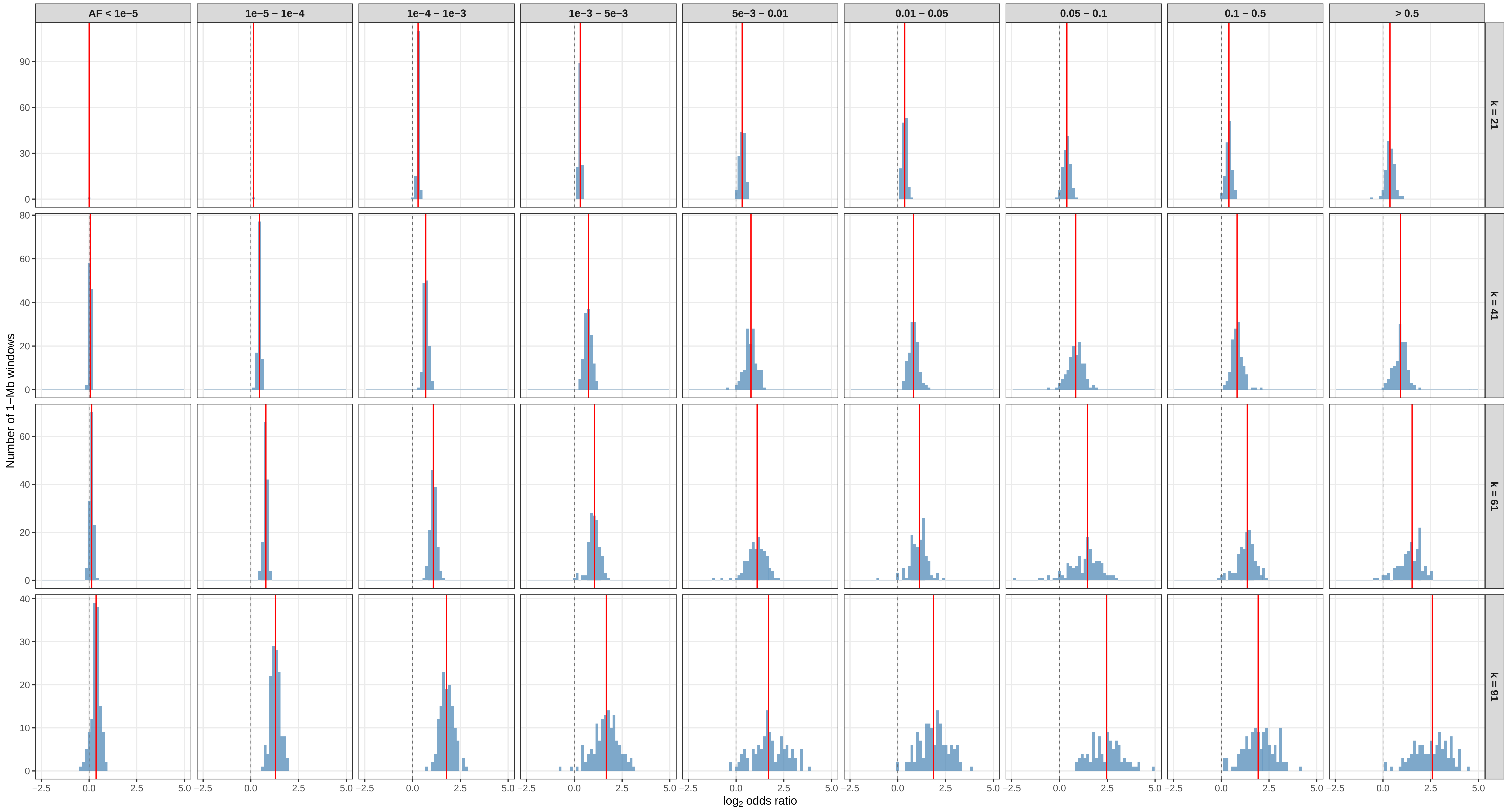

#### Chromosome 13

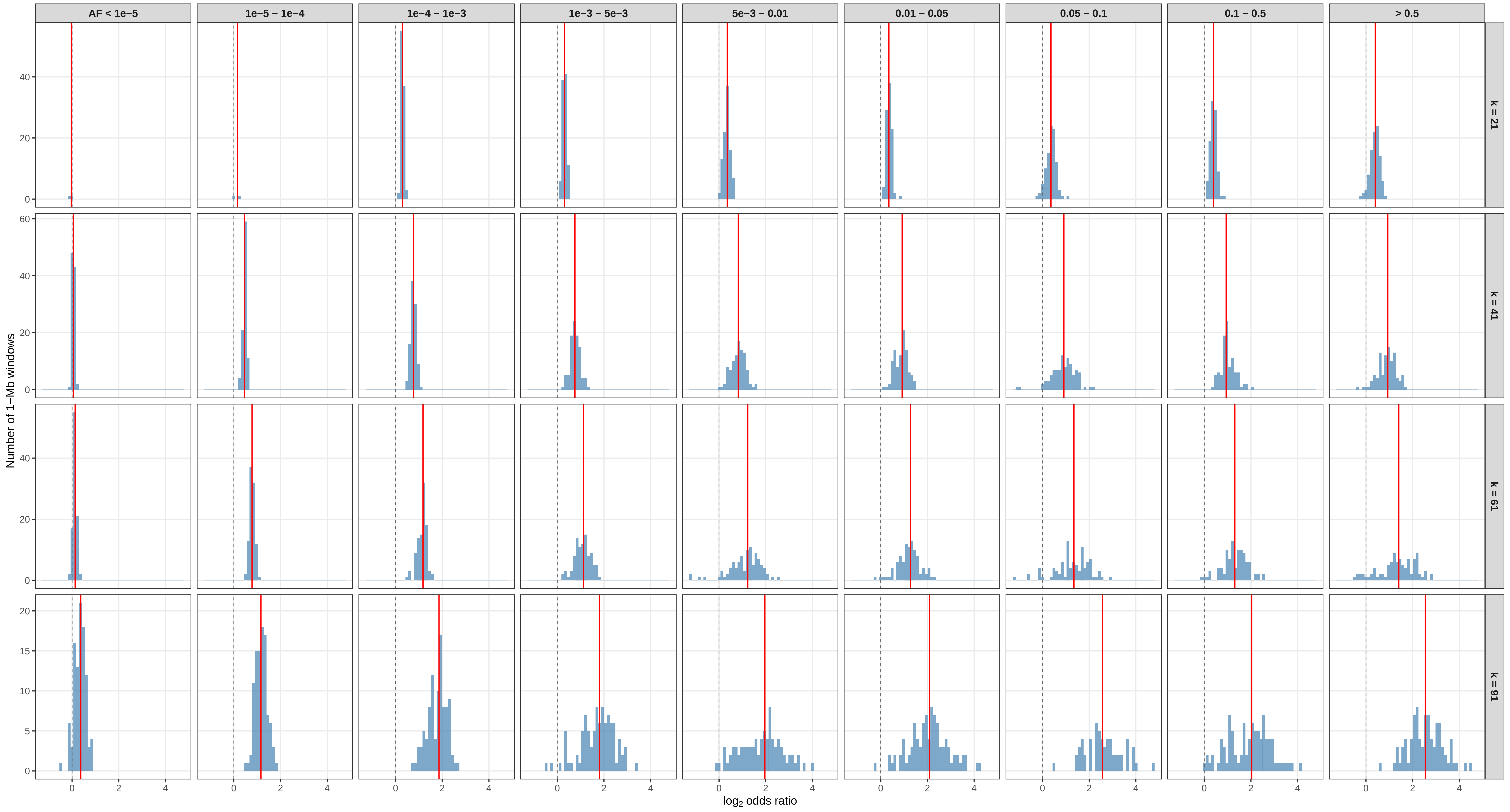

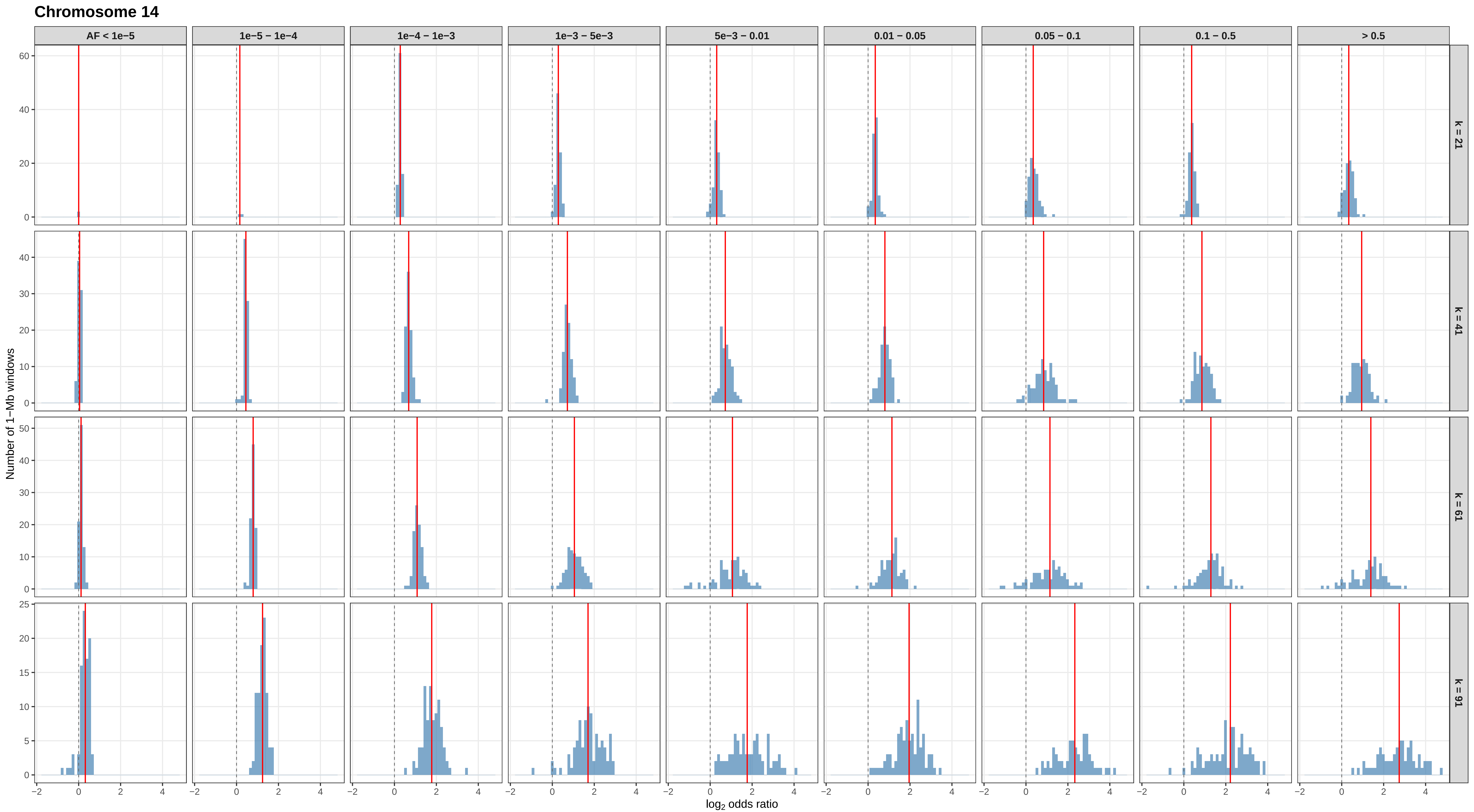
Chromosome 14
AF < 1e−5
1e−5 – 1e−4
1e−4 – 1e−3
1e−3 – 5e−3
5e−3 – 0.01
0.01 – 0.05
0.05 – 0.1
0.1 – 0.5
> 0.5
k = 21
k = 41
k = 61
k = 91
Number of 1−Mb windows
log2 odds ratio

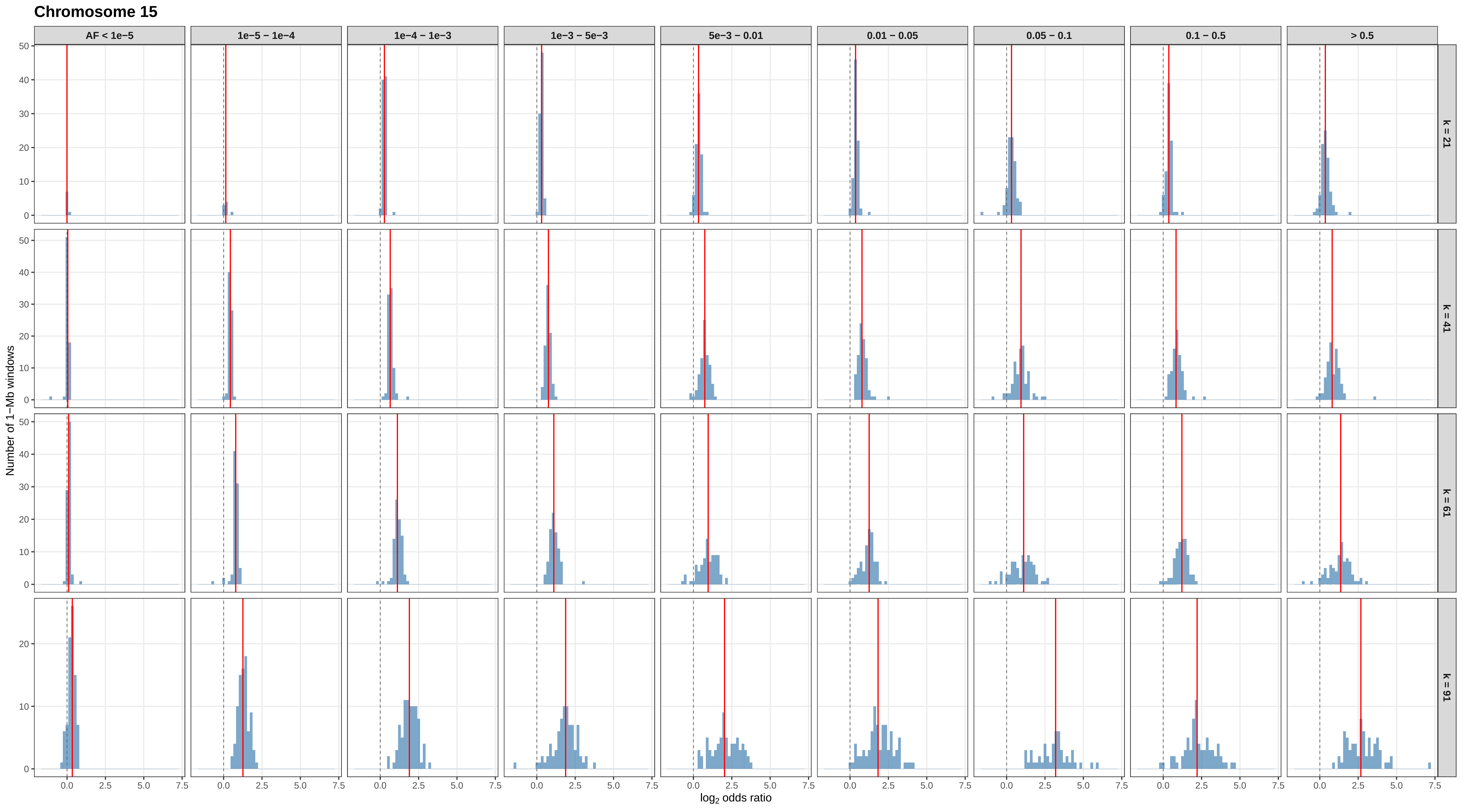

Chromosome 15
AF < 1e−5
1e−5 – 1e−4
1e−4 – 1e−3
1e−3 – 5e−3
5e−3 – 0.01
0.01 – 0.05
0.05 – 0.1
0.1 – 0.5
> 0.5
k = 21
k = 41
k = 61
k = 91
Number of 1−Mb windows
log2 odds ratio

#### Chromosome 16

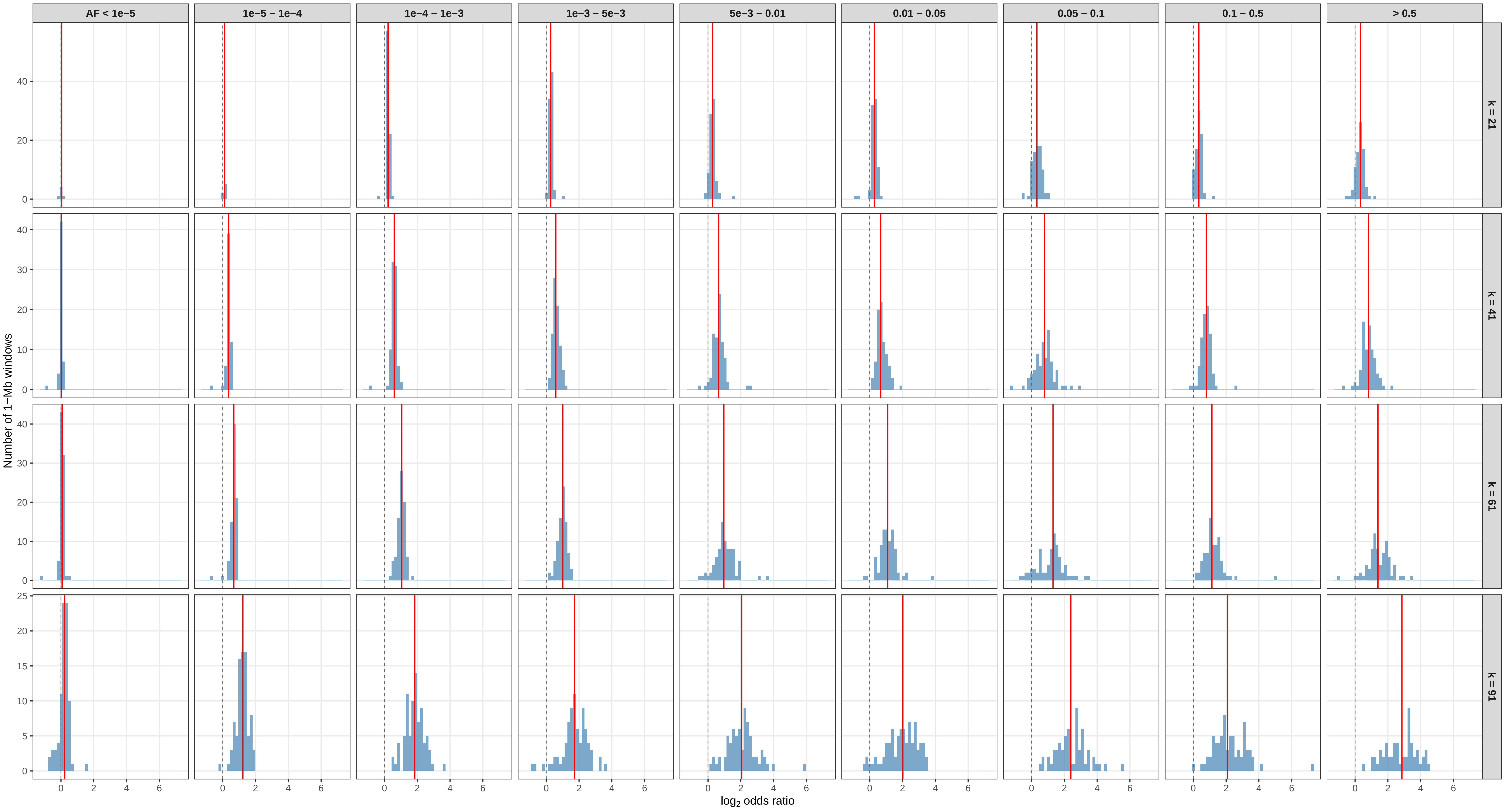

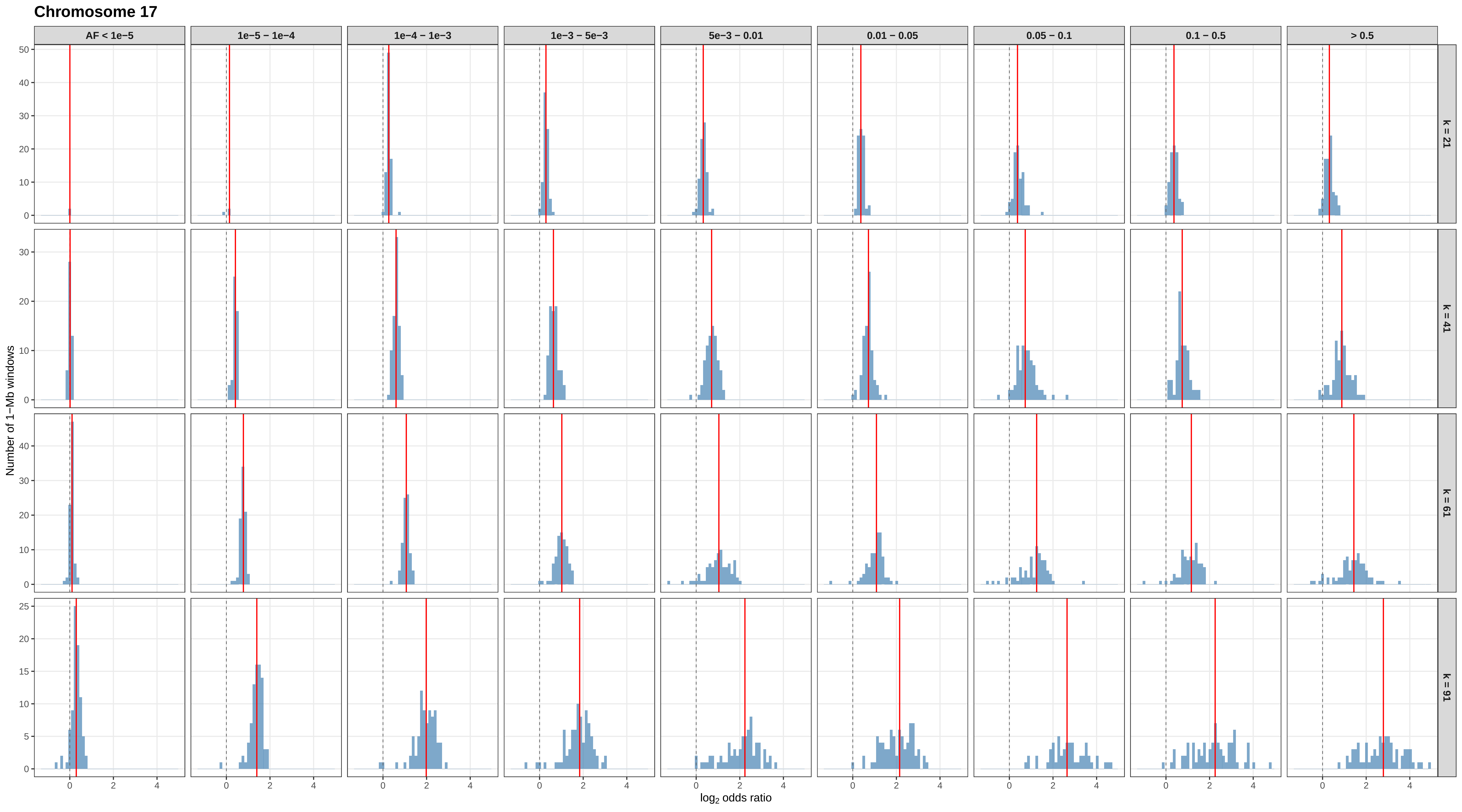
Chromosome 17
AF < 1e−5
1e−5 – 1e−4
1e−4 – 1e−3
1e−3 – 5e−3
5e−3 – 0.01
0.01 – 0.05
0.05 – 0.1
0.1 – 0.5
> 0.5
k = 21
k = 41
k = 61
k = 91
Number of 1−Mb windows
log2 odds ratio

#### Chromosome 18

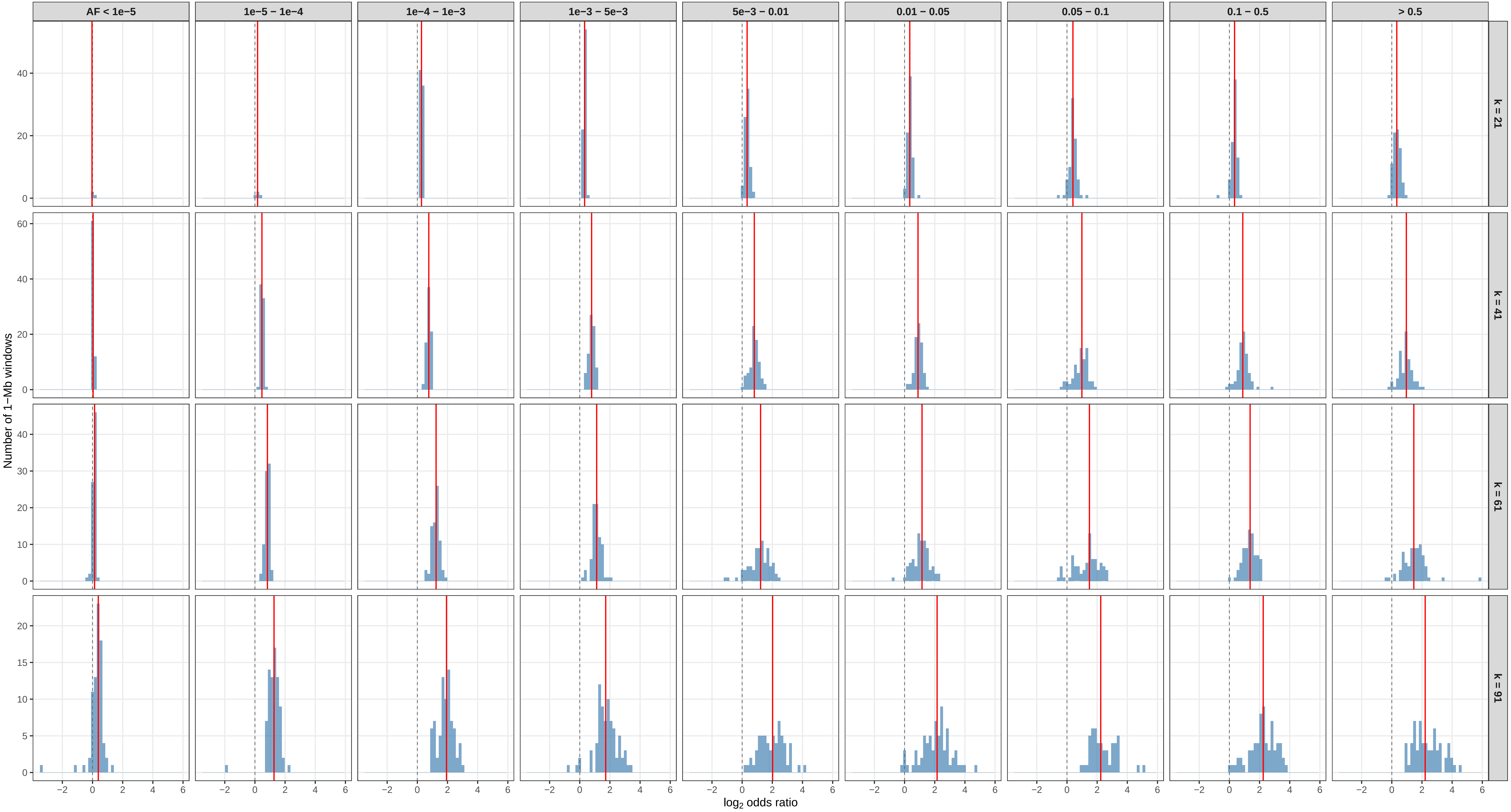

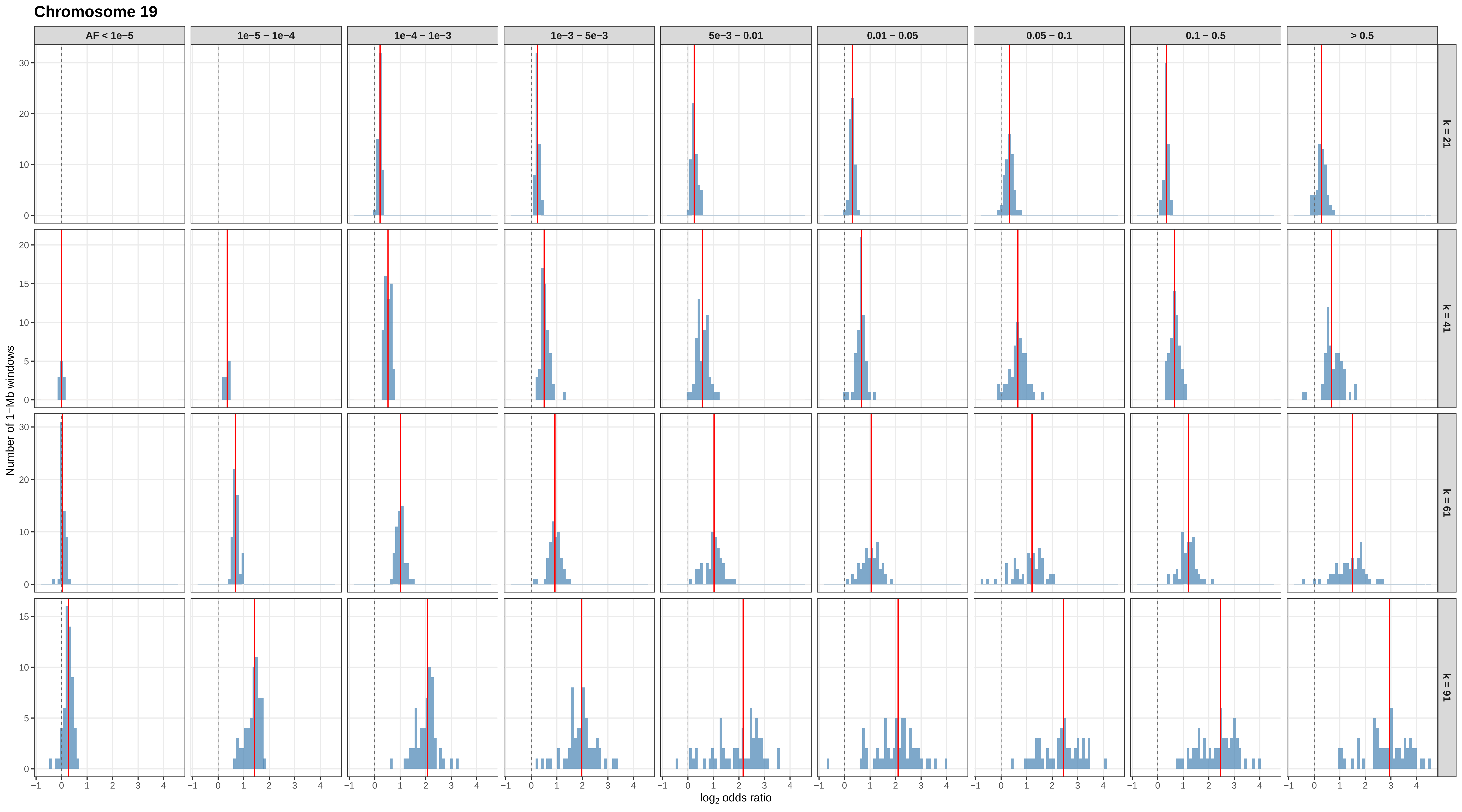

Chromosome 19
AF < 1e−5
1e−5 − 1e−4
1e−4 − 1e−3
1e−3 − 5e−3
5e−3 − 0.01
0.01 − 0.05
0.05 − 0.1
0.1 − 0.5
> 0.5
k = 21
k = 41
k = 61
k = 91
Number of 1−Mb windows
$\log_2$ odds ratio

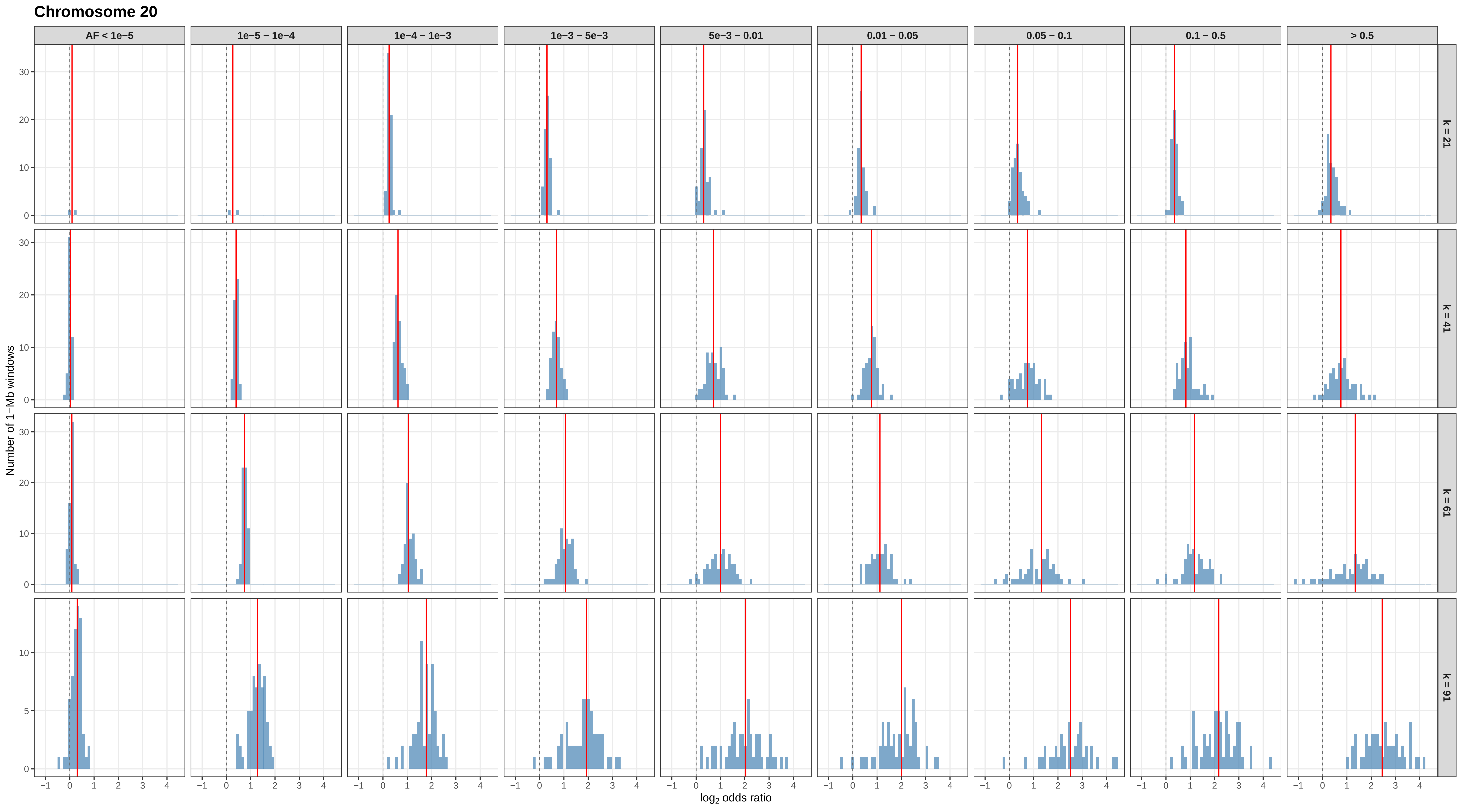

Chromosome 20
AF < 1e−5
1e−5 – 1e−4
1e−4 – 1e−3
1e−3 – 5e−3
5e−3 – 0.01
0.01 – 0.05
0.05 – 0.1
0.1 – 0.5
> 0.5
k = 21
k = 41
k = 61
k = 91
Number of 1−Mb windows
log2 odds ratio

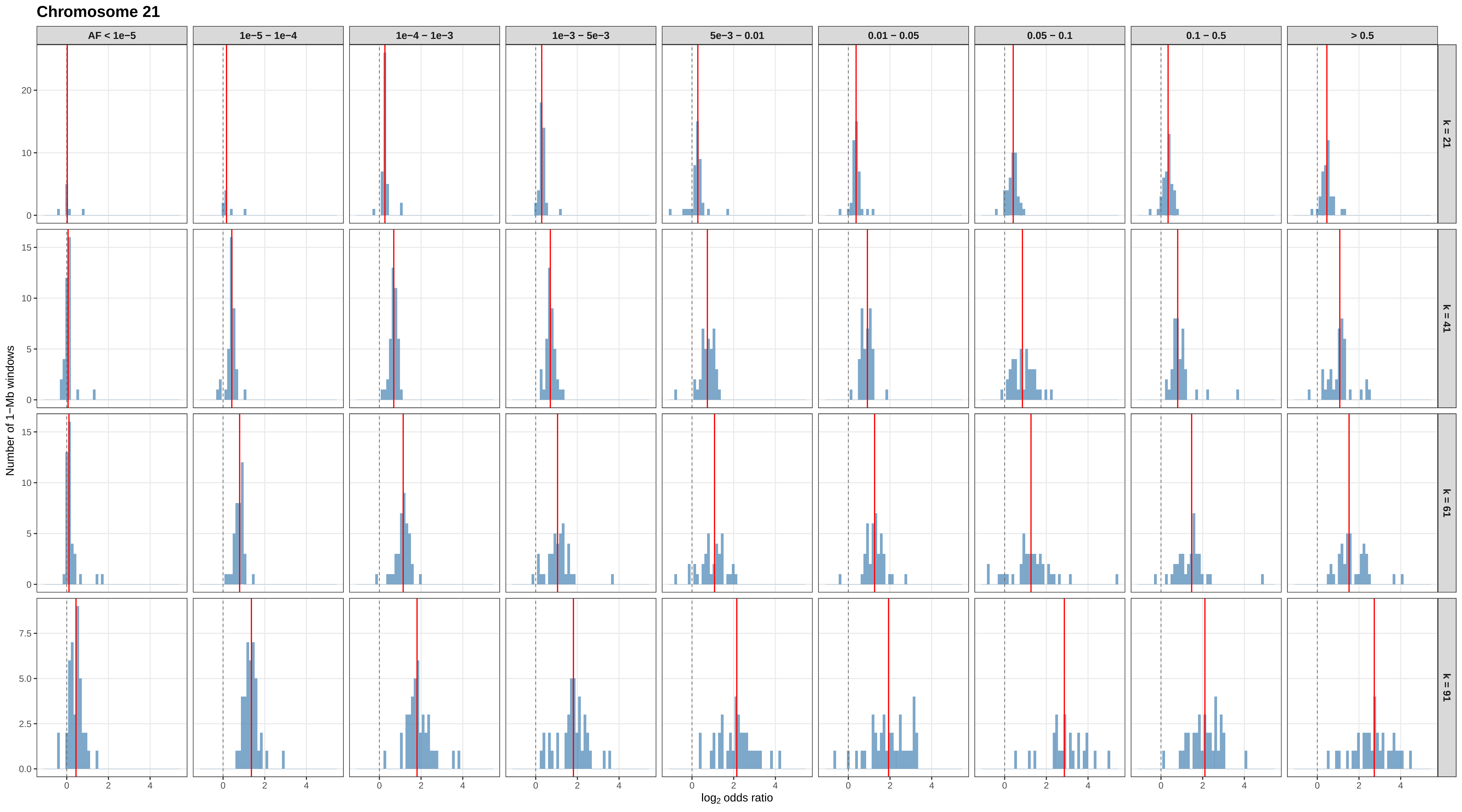

Chromosome 21
AF < 1e−5
1e−5 − 1e−4
1e−4 − 1e−3
1e−3 − 5e−3
5e−3 − 0.01
0.01 − 0.05
0.05 − 0.1
0.1 − 0.5
> 0.5
k = 21
k = 41
k = 61
k = 91
Number of 1−Mb windows
log2 odds ratio

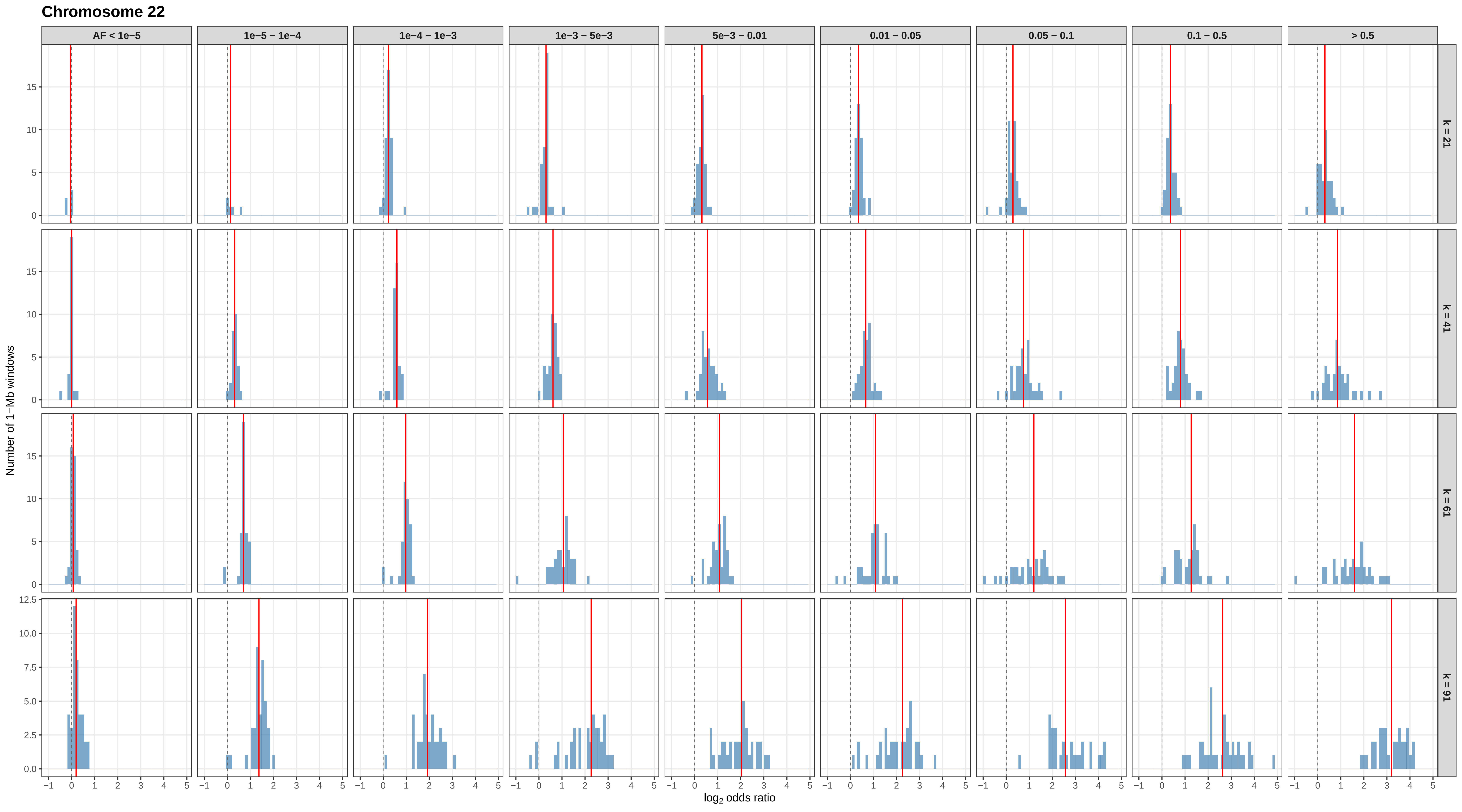
Chromosome 22
AF < 1e−5
1e−5 − 1e−4
1e−4 − 1e−3
1e−3 − 5e−3
5e−3 − 0.01
0.01 − 0.05
0.05 − 0.1
0.1 − 0.5
> 0.5
k = 21
k = 41
k = 61
k = 91
Number of 1−Mb windows
log2 odds ratio

#### Chromosome X

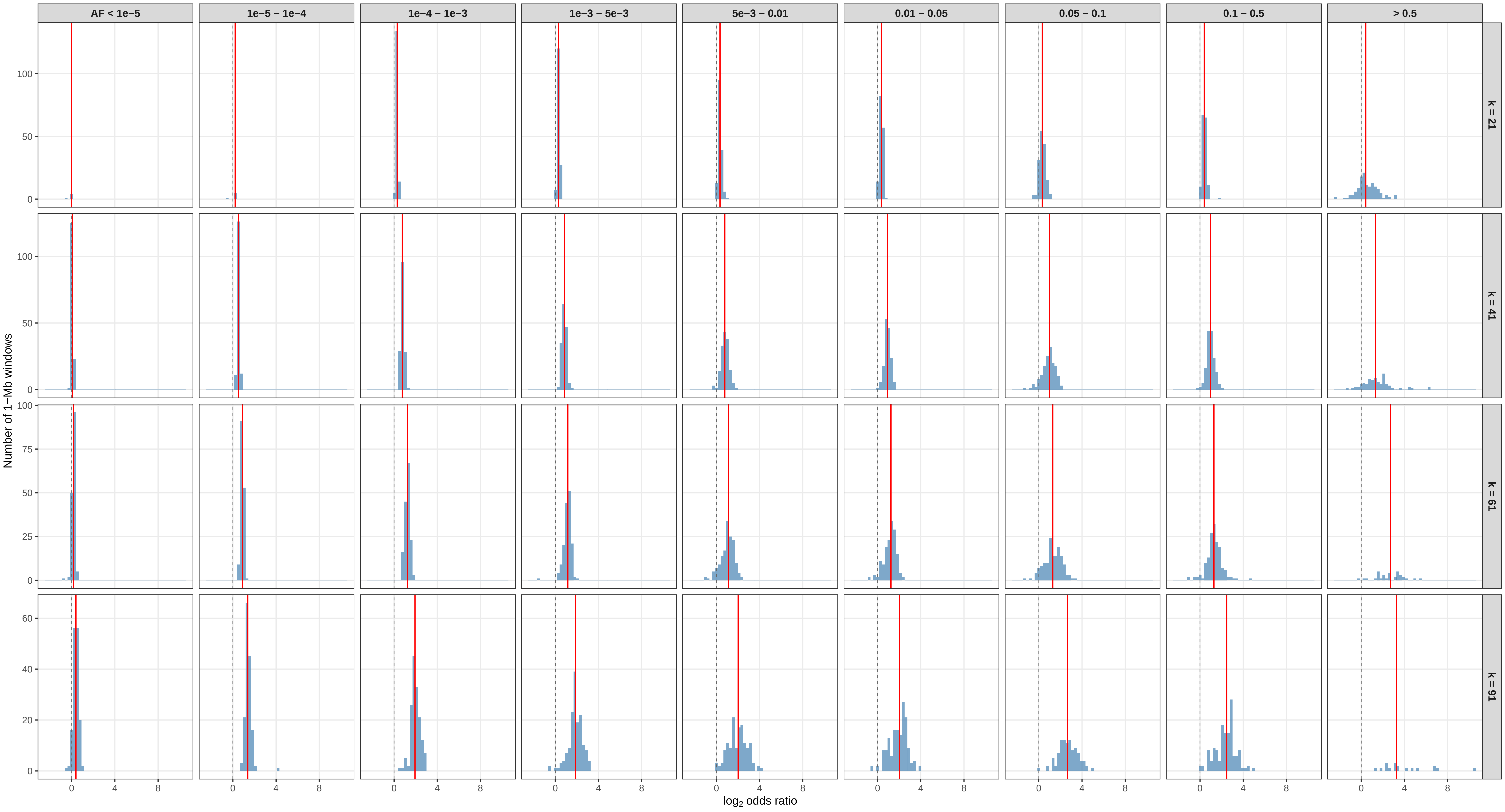

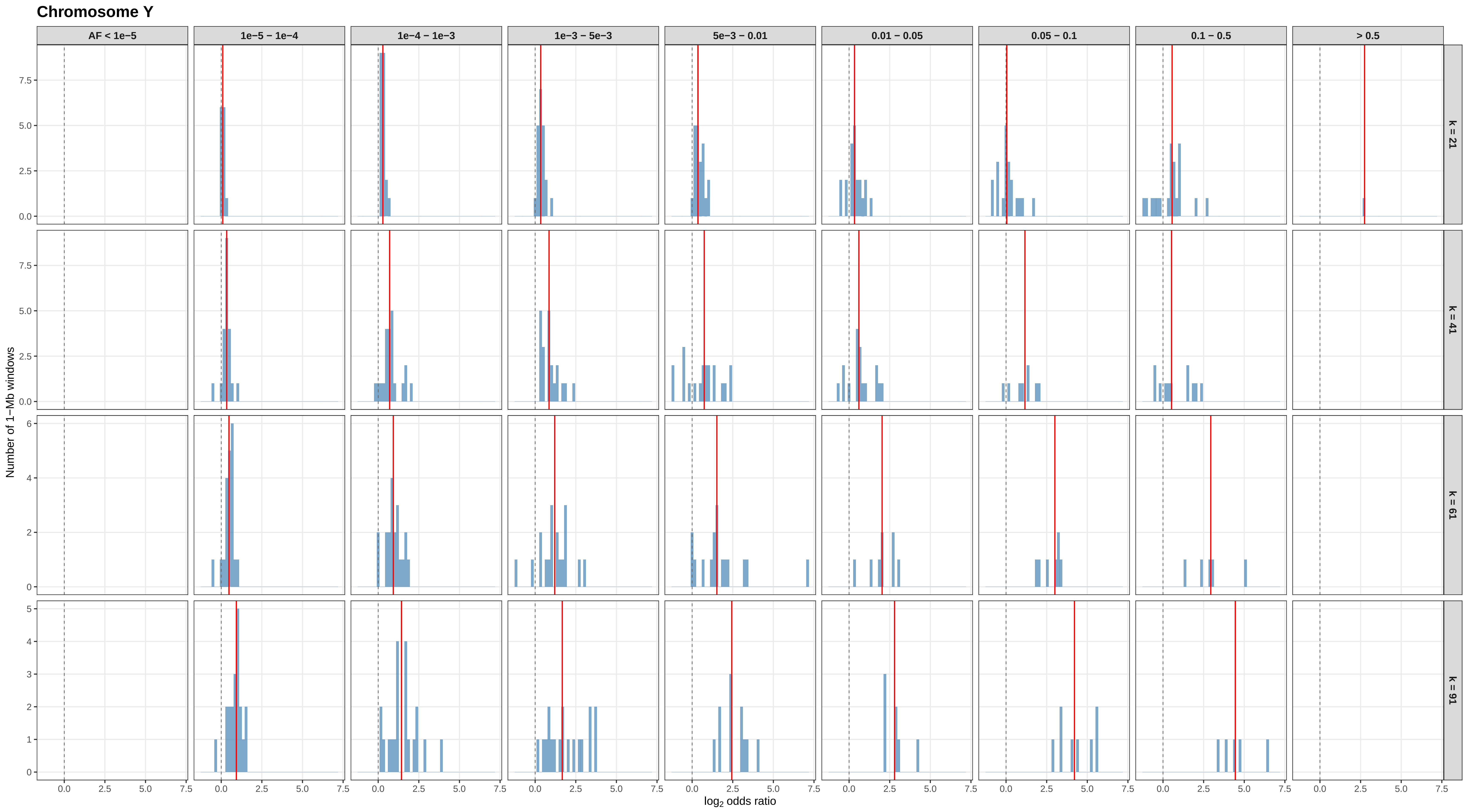

Chromosome Y
AF < 1e−5
1e−5 − 1e−4
1e−4 − 1e−3
1e−3 − 5e−3
5e−3 − 0.01
0.01 − 0.05
0.05 − 0.1
0.1 − 0.5
> 0.5
k = 21
k = 41
k = 61
k = 91
Number of 1−Mb windows
log2 odds ratio
0.0
2.5
5.0
7.5
0
2
4
6
1
3
5

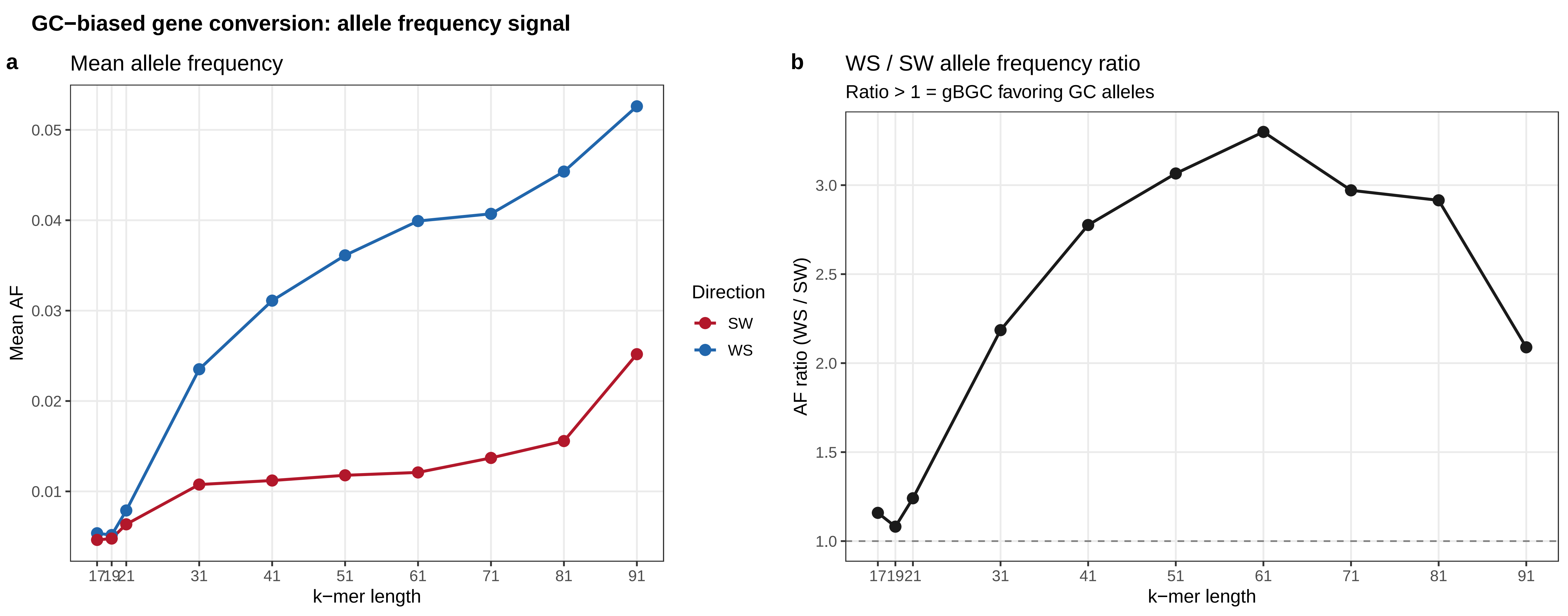
GC−biased gene conversion: allele frequency signal
a
Mean allele frequency
Mean AF
0.05
0.04
0.03
0.02
0.01
17
19
21
31
41
51
61
71
81
91
k−mer length
Direction
SW
WS
b
WS / SW allele frequency ratio
Ratio > 1 = gBGC favoring GC alleles
AF ratio (WS / SW)
3.0
2.5
2.0
1.5
1.0
17
19
21
31
41
51
61
71
81
91
k−mer length
